# Mapping the journey of a neuro-invasive virus in zebrafish larvae: contrasted viral spread and type I interferon responses in periphery and central nervous system

**DOI:** 10.1101/2024.05.19.594871

**Authors:** Valerio Laghi, Laurent Boucontet, Hannah Wiggett, Ingrid Colin, Payel Banerjee, Matthieu Simion, Ludovico Maggi, Sorana Ciura, Jérémie Guedj, Emma Colucci-Guyon, Jean-Pierre Levraud

**Affiliations:** Institut Pasteur, Unité Macrophages et Développement de l’Immunité, Centre National de la Recherche Scientifique (CNRS) UMR3738, Université Paris-Cité, 75015 Paris, France; Université Paris-Saclay, CNRS UMR9197, Institut Pasteur, Université Paris-Cité, Institut des Neurosciences Paris-Saclay, 91400 Saclay, France; TEFOR-Paris Saclay, Université Paris-Saclay, CNRS UAR2010, 91400 Saclay, France; Université Paris Descartes Hôpital Necker-Enfants Malades, Institut Imagine, 75015 Paris, France; Université de Paris, IAME, INSERM, 75018 Paris, France

## Abstract

Organ-to-organ dissemination of viruses is a critical feature of host-virus interactions. Neuroinvasive viruses can enter the central nervous systems (CNS), which often results in death or permanent neurological impairment. The complex mechanisms underpinning this spread are poorly understood, as they depend on a variety of parameters, including initial site of entry, route of access to the CNS, and immune responses. To better understand these phenomena, we analyzed the spatial dynamics of Sindbis virus (SINV) dissemination in the transparent zebrafish larva. Using fluorescent reporter viruses, we observed that SINV readily invaded the CNS after inoculation at various peripheral sites. Both sensory and motor neurons were gateways to the CNS, with 2 to 3 occurrences of CNS entry par larva. While peripheral infection was transient, CNS infection was persistent and more variable. Within the CNS, viral dissemination resulted both from long-distance axonal transport and short distance shedding. Type I interferon (IFN) response was systemic in the periphery, and local in the CNS; nevertheless, it played a key role in controlling the infection in both compartments, as shown by IFN receptor knockdown or with IFN triple mutants. We built a mathematical model of the infection based on our quantitative imaging, providing additional insight. The rate of new virion production was estimated around 1 to 2 infective virions per productively infected cell per hour. The model also found that IFN did not only prevent new infections but accelerated the death of infected cells, and had a stronger effect in periphery than CNS. This study paves the way for a deeper understanding of the mechanisms at play in pathogen- and host response-mediated neurological damage during viral encephalitis.

## Introduction

Invasion of the central nervous system (CNS) is one of the worst possible events during a viral infection (Swanson and McGavern 2015). This remains relatively rare because the CNS is protected by specialized barriers, notably the blood-brain barrier (BBB). When it occurs, both the direct viral cytopathic effect and the inflammatory response induced may cause serious damage, often resulting in death or permanent neurological impairment (Venkatesan 2015). The complex interplay between the virus and the host response has been studied in a variety of animal models, but their dynamics remain poorly understood, largely because of the difficulty of following these events in the CNS. Mathematical modeling has provided insights in viral infection dynamics in the periphery based on repeated blood sampling (Perelson 2002), but longitudinal data were not available for the CNS.

The larva of the zebrafish Danio rerio has recently emerged as a powerful model to study host-pathogen interactions. Its small size and transparency are key advantages; by full-body intravital imaging, using pathogens encoding fluorescent reporter genes, it is possible to follow their organ-to-organ dissemination in real time and at high resolution (Tobin, May, and Wheeler 2012). Here, we took advantage of these properties to understand the propagation of a neuroinvasive virus in vivo.

Sindbis virus (SINV) is a single-strand positive RNA virus belonging to genus *Alphavirus*, transmitted by mosquitoes to its natural bird hosts but also sometimes to mammals including humans. While SINV causes only mild symptoms in humans (Adouchief et al. 2016), other members of this genus include major human pathogens such as chikungunya virus (CHIKV) (Strauss and Strauss 1994a). Several members of this group, such as Eastern Equine Encephalitis Virus and Venezuelan Equine Encephalitis Virus are known to cause fatal encephalitis in humans. While CHIKV was mostly reputed to cause arthralgia and myalgia, the massive breakthrough that occurred in La Reunion Island revealed that it was also encephalitogenic, particularly in newborns (Das et al. 2010). SINV is used to study viral encephalitis in mice. It is generally injected directly in the brain, but can propagate from the periphery to the CNS in newborns, particularly with strains adapted by multiple intracerebral passages (Lustig et al. 1988). More generally, SINV is well known to infect neurons and replication-deficient strains are used as a tool to transduce neurons and trace neural circuits (Furuta et al. 2001).

We have previously established that SINV was neuroinvasive in zebrafish larvae, relying on axonal transport, but not on BBB infection or opening, or on macrophage-mediated entry, to gain access to the CNS after intravenous (IV) inoculation (Passoni et al. 2017). SINV induced a rapid and strong type I interferon (IFN) response in zebrafish larvae after IV injection; knocking down IFN receptors revealed that this response was protective, preventing rapid death of infected larvae (Boucontet et al. 2018). Nevertheless, we observed that in wild-type larvae, SINV infection persisted or kept progressing in the CNS, in contrast to periphery where it appeared to be transient. We had previously reported similar kinetics of IFN induction, and comparable persistence specifically in the CNS, in zebrafish infected with CHIKV (Palha et al. 2013), despite a different CNS entry route (Passoni et al. 2017). This led us to hypothesize that localization of IFN responses may play a key role in organ-specific persistence of CHIKV (Levraud et al. 2014).

To address these questions, we performed detailed and quantitative analyses of SINV propagation from periphery to CNS. Because IV injections result in highly variable infection patterns, presumably due to stochastic initial infection of a few cells among a very large target pool accessible from the bloodstream, we tested other injection routes which also systematically resulted in CNS entry, but with more predictable patterns. Focusing on intramuscular (IM) injections in caudal somites, we observed that infection of peripheral cells was transient, while CNS infection was progressive and more variable. We identified sensory neurons in dorsal root ganglia (DRG) as a major but not exclusive entry way to the spinal cord and observed short- and long-range propagation modes within the CNS. Knocking down IFN receptors resulted in much stronger infection, most notably in periphery where transiency was lost, but also in the CNS; similar results were observed with type I IFN-deficient mutant larvae. Mathematical modeling indicated that strong differences in IFN responsiveness must exist between periphery and CNS to account for these observations, which was consistent with imaging of IFN reporter zebrafish larvae. This study paves the way for a deeper understanding of the mechanisms at play in pathogen- and host response-mediated neurological damage during viral encephalitis.

## Results

### Organ to organ propagation of SINV

To establish a global picture of SINV dissemination in zebrafish larvae, we compared different routes of virus inoculation. We co-injected two viruses encoding eGFP or mCherry with an otherwise identical backbone derived from the pTE3’2J SINV clone, previously established to be a of relatively low virulence in zebrafish (Boucontet et al. 2018; Passoni et al. 2017). The virus mix was injected to 3 days post fertilization larvae (dpf), either intravenously (IV), intramuscularly (IM), inside the pericardial cavity, or in the spinal cord (Figure 1A). Intracerebral injections were also tested but not pursued because of massive brain infection and rapid mortality. For each route of injection, two dozen larvae were followed, split over two independent experiments. Fluorescence images of infected larvae were taken daily up to 4 days post injection (dpi) using a widefield microscope at low magnification, allowing us to follow the global infection pattern of each larva over time (Figure 1B).

**Figure 1:**
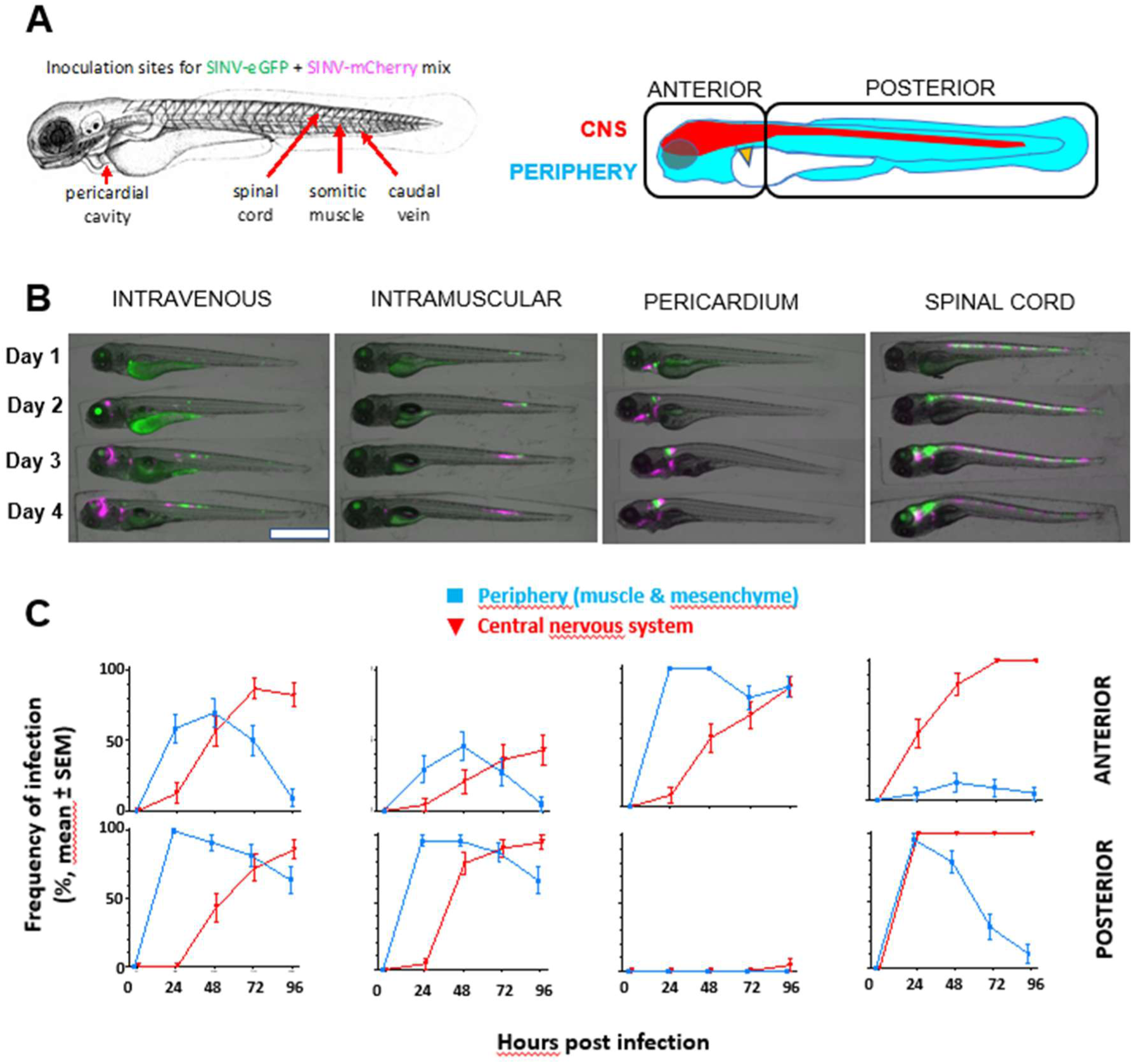
Dynamics of infection associated to different injection routes. **A)** Scheme of 3dpf zebrafish larva showing routes of injection (LEFT) and compartmentalization of larva for quantification of viral presence frequency patterns inside (red) or outside (cyan) the CNS in anterior or posterior part of the larva (RIGHT). “PC” Pericardial Cavity, “IM” Intramuscular, “IV” Intravenous and “SC” Spinal Cord. **B)** Representative images of larvae injected in different sites with a mix of SinV-GFP (green) and SinV-mCherry (magenta) strains, and followed for 4 days upon injection. **C)** Quantification of viral presence frequency patterns relative to different routes of injection divided by anterior (head)/posterior (tail) area and periphery (cyan)/CNS (red) compartments (n=24; 2 independent experiments). Scale bar 1mm.

Images were scored blindly to determine infected organs with either the eGFP or the mCherry virus (Table S1). To verify that the two viruses are equally infectious, we compared the localization of mCherry and eGFP foci. We did not notice an obvious bias except in the proximal kidney tubule, which we know results from reuptake of fluorescent proteins from the renal filtrate, which are then directed to acidified endosomes in tubule cells (Eshbach and Weisz 2017) where eGFP but not mCherry fluorescence is quenched; we also routinely observe a comparable signal in uninfected mCherry-expressing transgenic larvae. Thus, this difference in the kidney does not result from infection of this organ. We used Fisher’s exact test to compare the frequencies of infection by the GFP and mCherry virus in 5 areas for the 4 injection sites and 4 time points (i.e., 80 comparisons) (Figure S1). We found only 3 cases where the test indicated significantly different frequencies (p<0.05); with 80 tests performed, this is what is expected from random fluctuations. Therefore, as expected, the two isogenic viral clones display similar tropism and infection dynamics.

To compare the infection patterns over time and inoculation route, we tabulated the frequency of infection of 36 different body areas, deduced from our low-resolution images (Table S1) and performed principal component analysis (PCA) of this dataset (Figure S2A). The two independent replicate groups at a given injection site behaved similarly, indicating that infection patterns were reproducible. Spinal cord injected animals were quite separate from other samples. Interestingly, all groups showed a similar trend towards a higher PC1 coordinate value with time, with peripherally injected animals progressively getting closer to spinal cord injected animals. Accordingly, the sites with the highest positive weights in PC1 were the brain and spinal cord regions (Figure S2B). Thus, this evolution in the PCA plane over time essentially reflects the progressive invasion of the central nervous system by this neuroinvasive virus.

Bloodstream injection (Figure 1B, first row) yielded the broadest pattern of early infection. Tail muscle was often infected, undoubtedly because of the needle having to pass through somite to reach the main tail blood vessels. This in reflected by the close trajectories of IV and IM groups on the PCA graph (Figure S2A). Positive organs distant from the injection site, and thus likely infected by bloodborne virus, included the liver, jaw, gills, heart, peripheral nerve ganglia, and the conspicuous syncytial yolk cell. Infection of these distant sites was quite variable from larva to larva. Consistent with our initial description of the SINV model in zebrafish (Passoni et al. 2017), infection propagated later to the central nervous system, where it lasted, unlike peripheral infection which was mostly transient. We plotted the frequencies of infection of significant areas of peripheral tissue and CNS in the anterior and posterior regions (Figure 1C) and this trend is observed in both halves.

Intramuscular injections (Figure 1B, second row) resulted in a more localized pattern. Early injection was typically restricted to the injected somite, then propagated to the closest somite (but rarely beyond) and often to nearby fin mesenchyme. Spinal cord infection almost systematically ensued, first at the position slightly anterior to the injected somite, then propagating both anteriorly and posteriorly. Occasional infections were observed in the anterior regions (Figure 1C) presumably because of some leakage of the inoculum into the blood when injecting IM.

Pericardial cavity injections (Figure 1B, third row) also resulted in a reproducible pattern. Heart, gill, and jaw mesenchyme were infected early, as well as mandibular muscles. Infection then propagated to the hindbrain. Remarkably, infection was not observed in the posterior half (Figure 1C).

Spinal cord injections (Figure 1B, fourth row) resulted in early infection of entire segments of the spinal cord, which then propagated to the rest of the spinal cord, hindbrain, and rest of the brain. Some tail muscle infection was also observed at early times; we interpret this as a primary event because the needle had to pass through muscle to reach the spinal cord. By contrast, almost no infection of anterior peripheral regions was observed (Figure 1C).

A striking mutual exclusion of the eGFP and mCherry positive cell patches was observed in spinal cord injected larvae (Figure 1B). On closer examination, it was also systematically observed at other sites with all injections. The only cell for which we observed unquestionable co-infection by SINV-GFP and SINV-mCherry was the syncitial yolk cell (in 6/24 IV-injected and 5/24 IM-injected larvae).

Infections near the injection site almost always included both eGFP and mCherry positive cells; by contrast, infected patches appearing at a distant site (except yolk) were typically single-colored, indicating initial seeding by a single virion. A bottleneck effect was also observed for neuro-invasion events: among 20 IM-injected larvae with clear dual color initial muscle infection, 12 later displayed dual-color and 7 single-color invasion of the spinal cord; while among 24 pericardial cavity-injected larvae with dual color infection of the jaw/gill area, 7 later had a dual-color infected brain and 15 a single-color infected brain. Assuming that multiple instances of BBB crossing occur independently from each other, we calculated, using Poisson’s law (Supplementary text 1), that there are ∼3 events of successful CNS entry (range, 1.7 to 4.4) after IM injection and ∼2 such events (range, 1.2 to 2.6) after pericardial injection. In the tail, infection of caudal muscles preceded invasion of spinal cord, while anteriorly, infection of facial/jaw muscles preceded hindbrain invasion, which was consistent with the axonal transport of SINV we had established previously (Passoni et al. 2017). For the rest of the study, we decided to focus on IM tail injections because of its clear pattern of muscle to spinal cord propagation, relative thinness of the infected region facilitating imaging, and relevance of entry route for this mosquito-transmitted virus.

### Sensory and motor neurons contribute to spinal cord invasion

In our earlier work (Passoni 2017), we had already noted that infection of muscle fibers was followed by infection of neurons that innervate this somite. Here, we analyzed these events in detail by inoculating SINV-mCherry to Tg(*elavl3*:GFP) (also known as *HuC*:GFP) reporter zebrafish, labelling all neurons, or Tg(*vsx2*:GFP) (also known as *axl1*:GFP), labelling a subset of interneurons. Time-lapse imaging by confocal microscopy detected the first infected cells at ∼7 hours post-injection (pi) (6.86±0.38 hours, mean±SD, n=7 larvae). During the first 24 hours, infection progressed in the periphery (Figure 2A; movie S1); infected cells prominently included muscle fibers but also numerous stromal cells between myotomes (in myoseptum and myocommata), in the caudal hematopoietic areas, in the conjunctive tissue dorsal to the somites (Figure 2A, movie S1), and sometimes in fin mesenchyme (Figure 2B, movie S2).

**Figure 2:**
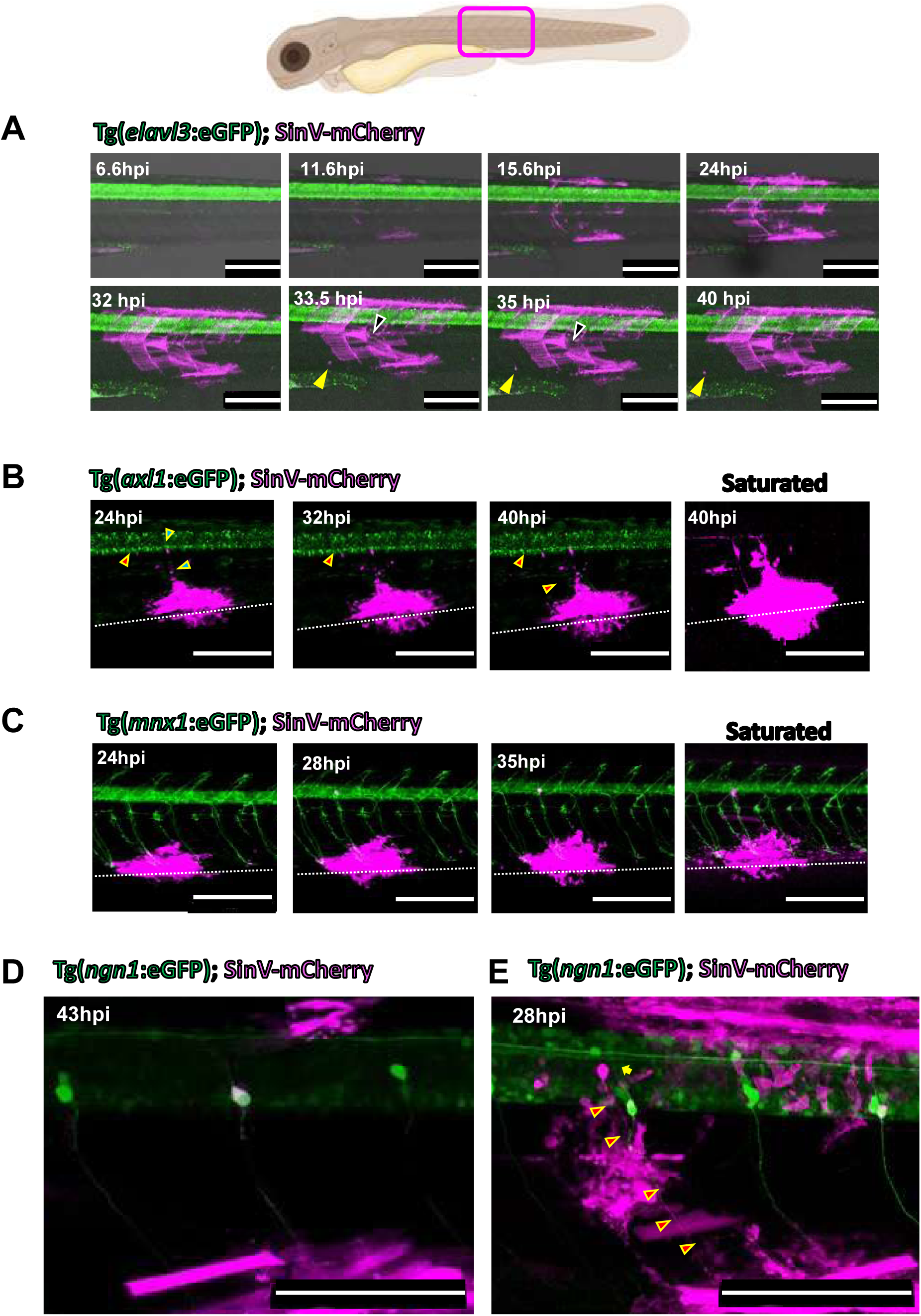
Timelapse imaging after IM inoculation of SINV, showing infection of the somite and identification of the first infected neurons. Confocal in vivo imaging of zebrafish larvae expressing GFP transgenes (green) infected IM with SINV:mCherry (magenta), maximal projections. **A)** Selected time-points from timelapse of Tg(elav3:GFP) larvae showing somite infection developing during the first and second days. Note that top and bottom rows correspond to two different larvae. Yellow arrowheads point to a wandering leucocyte containing fluorescent material. Black arrow points to a dying muscle fiber. **B)** Selected time-points from in vivo timelapse of a Tg(axl1:GFP) larva showing infected neurons innervating the infected zone in the ventral somite. Green-filled arrowheads point to an infected neuron with its axon already labelled by mCherry at start of timelapse; red-filled arrowhead point to another neuron with its axon becoming visibly labelled only later. The dotted line indicates the boundary between the trunk and the ventral fin. **C)** Selected time-points from in vivo timelapse of an infected Tg(mnx1:GFP) larva, revealing that the first infected neuron is not a motoneuron as its axon is not labelled by GFP. **D,E)** Confocal images of infected Tg(ngn1:GFP), showing infection of a DRG sensory neuron (D) and of a motor neuron, the axon of which is shown by red-filled arrowheads (E). Scale Bar, 200µm in A-C, 100µm in D-E.

During the second day pi, infection largely stalled in the periphery (Figure 2A, bottom row; movie S3). Interestingly, movement of bright mCherry specks was often observed, suggesting that fragments of dead infected cells had been phagocytosed by a wandering leukocyte (movie S3). By contrast, infection started to be visible in the spinal cord. When a neuron got infected, we first observed fluorescence in its soma, and in its axon a few hours later, as expected for a soluble fluorescent protein produced in the soma. The first visibly virus-fluorescent axon invariably innervated the infected muscle area, suggesting that this axon had carried the virus to this neuron soma a few hours earlier (Figures 2B, C).

Somites are innervated by motor and sensory neurons. To identify which neuron subtype was infected first, we performed time-lapse imaging of the infection using the motoneuron reporter line Tg(*mnx1*:GFP) infected with SINV-mCherry. We could identify the first neurons becoming infected, first by their soma becoming mCherry-positive and a few hours later by their axons being labelled. Figure 2C and movie S4 show infection of neuron that became just visible at 24 hpi, at the start of the timelapse. Three hours later, its axon was visibly labelled; as expected, it innervated the infected ventral trunk area. This axon ran close but distinct from that of the GFP+ motor neurons, and 3D reconstruction (movie S5) indicated that the soma of this axon was located lateral to the spinal cord, in a dorsal root ganglion (DRG). Thus, in this case, the first infected neuron was a sensory DRG neuron.

To directly visualize infection of sensory DRG neurons, we inoculated SINV-mCherry into Tg(*ngn1*:GFP) fish, in which these cells are strongly labelled. We confirmed that a DRG sensory neuron often became infected before spinal neurons (Figure 2D; movie S6). In at least one case, however, when starting the time-lapse at 25 hpi, the most heavily infected neuron was clearly a motoneuron (Figure 2E), movie S7); a DRG sensory neuron was also infected, but its axon, unlike the axon of the infected motoneuron, was not mCherry-positive, indicating it had been infected later. Thus, both sensory and motor neurons may be the first infected neuron mediating SINV entry to the CNS.

Overall, our data suggest that CNS entry is slightly more frequent via sensory neurons than via motoneurons; indeed, from our timelapse movie series, we observed that, at 24 hpi, one or more DRG neuron was already infected in 6 out of 8 fish, while one or more motoneuron was infected in 4 out of 8 fish. Considering that our statistical analysis from 2-color virus IM injections revealed 3 independent CNS entry events on average, entry in the spinal cord may be mediated by both muscle-innervating neuronal subtypes in a given fish.

### Short- and long-range dissemination of SINV within the spinal cord

We then examined how the infection propagates inside the CNS. We hypothesized that the virus could propagate either to its direct neighbors after budding from the soma or to distant cells using axon-mediated travel. Our high-resolution time-lapses revealed both short- and long-distance dissemination in the spinal cord. In one case (Figure 3A, Movie S8) starting at 24 hpi we initially see an infected motoneuron (magenta arrow) and, 50µm rostrally, three infected neurons close to each other (green arrow). In a few hours, we can detect a group of newly infected neurons surrounding this latter cluster (yellow arrows), as well as an isolated neuron 100µm upstream (cyan arrow), with a few immediate neighbors becoming fluorescent later (blue arrow). By 38hpi, axons that connect the distant clusters became visibly fluorescent.

**Figure 3:**
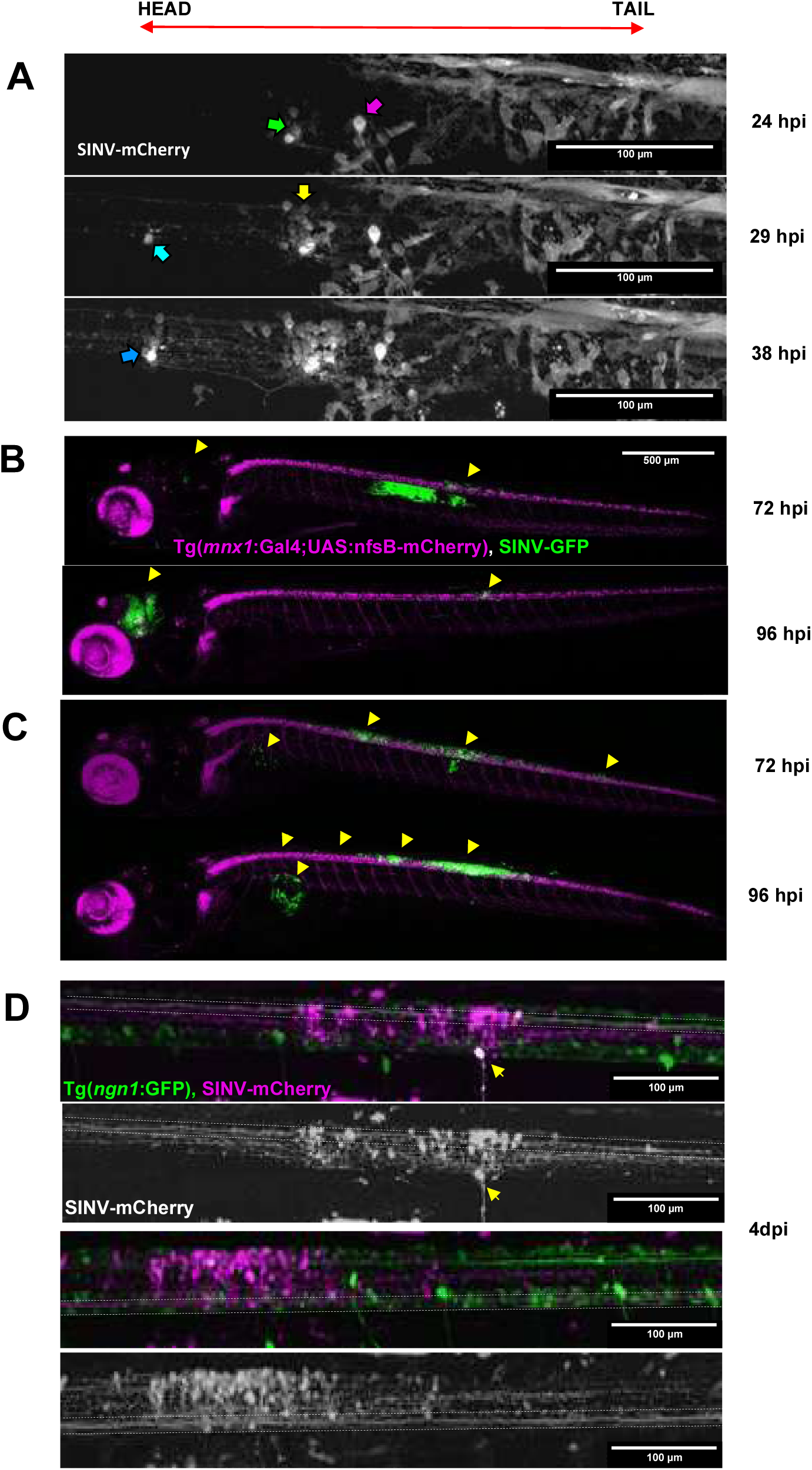
Propagation of SINV within the CNS. **A**. Progression of SINV infection in the spinal cord, confocal timelapse, maximal projection. (40x objective) Tg(ngn1:GFP) (GFP not shown, see movie S7 for uncropped two-color images) infected IM with SINV-mCherry (shown in grayscales). Neurons already infected at onset of the timelapse are shown with a magenta arrow (motoneuron) and green arrow (three neurons inside the cord). As infection progresses, a distant rostral neuron becomes infected (cyan arrow), while more neurons also become infected in the close vicinity of previous ones (yellow and blue arrows). **B,C)** Time points from confocal acquisition (10x objective) of Tg(mnx1:gal4; UAS:nfsB-mCherry), in magenta, infected with SINV-GFP (green). Arrows indicate infected cells: B) Long distance dissemination of SINV in absence of intermediate clusters between first point of entry in the spinal cord and brain secondary infection C) Short distance dissemination of SINV by clustering from primary point of entry in the spinal cord and secondary cluster in direction of the brain. **D)** Confocal images (40x objective) of infected long somatosensory axons connecting brain and periphery of Tg(ngn1:GFP) infected IM with 30 PFU SINV-mCherry (magenta). The yellow arrow indicates an infected DRG neuron and dotted line delimit respectively dorsal somatosensory long axons (Top) and ventral somatosensory long axons (Bottom). Black and white images show only the SINV-mCherry channel. Between dotted lines: infection of dorsal (top) and ventral (bottom) long somatosensory axon bundle.

Secondarily infected neurons were observed in all zones of the spinal cord without an obvious preference, including some *axl1*+ interneurons (not shown).

This axon-mediated propagation could span very long distances, as shown on fig 3B, where we see infected cells in the hindbrain between 82 and 96hpi, more than one millimeter away from the entry site in the spinal cord. Such a propagation to the brain was however not systematic; the larva shown in 3C, injected on the same day as the one in 3B, had a stronger and locally increasing spinal cord infection, but it did not propagate to the brain.

Long spinal axons labelled by the virus reporter could be detected either in the dorsal or the ventral spine (Figure 3D), suggesting that long distance propagation is mediated either by sensory or motor pathways.

### Death of infected sensory neurons

Although we imaged hundreds of infected spinal neurons in our time-lapse experiments, their death was very rarely seen. By contrast, death of infected DRG sensory neurons could be readily observed during timelapse imaging of Tg(ngn1:GFP) larvae infected with SINV-mCherry; Movie S9 provides an example of such an event, with the rostral-most DRG neuron in the imaged field already infected at the start of the movie (24hpi) and disappearing between 28:21 and 28:50 pi. Interesting, we can also observe a GFP-positive dorsal spinal axon decorated with double-labeled speckles, upstream and 120µm downstream of this DRG neuron, which we initially suspected to be local axonal bulges (possibly synapses) of this neuron, consistent with the typical T-shape of the spinal axon tract of DRG sensory neurons (Bernhardt et al. 1990). Indeed, a subset of these speckles disappear synchronously with the DRG cell soma. The other speckles all disappear together one hour later, suggesting they belonged to another infected DRG neuron, invisible because localized on the contralateral side, that died at around 30 hpi.

In addition, since DRG neurons have a stereotypical distribution and are easy to count in the Tg(*ngn1*:GFP) transgenic fish, we observed that when imaging of IM injected fish was started at 48 hpi or later, some DRG neurons were usually already missing at the level of the initial injection (1.57±0.90 at 72 hpi, n=7). Thus, infected DRG neurons appear to die a few hours after being visibly infected, much faster than spinal neurons.

### Entry via sensory neurons promotes brain infection

To functionally test for the relative role of motor and sensory neurons in spinal cord invasion, we performed depletion experiments (Figure 4). DRG neurons were prevented from differentiating from neural crest by injecting in eggs an antisense morpholino targeting the *erbb3b* transcripts (Dooley et al. 2013) (Figure 4A-D). Motor neurons were transiently depleted by a chemogenetic approach, treating Tg(*mnx1*:gal4 ; *UAS*:nfsB-mCherry) larvae with metronidazole (MZ) prior to infection (Davison et al. 2007) (Figure 4E-H). After infection by SINV, spinal cord was invaded in all groups (Fig 4B,F), consistent with our observation that both sensory and motor neurons may carry the virus from periphery to CNS. One notable difference, however, was that propagation of the infection to the brain was rarer in DRG-depleted animals, while its frequency was unchanged in motoneuron-depleted ones (Figure 4CG). Thus, the sensory neuron route appears to provide a privileged way to reach the hindbrain.

**Figure 4:**
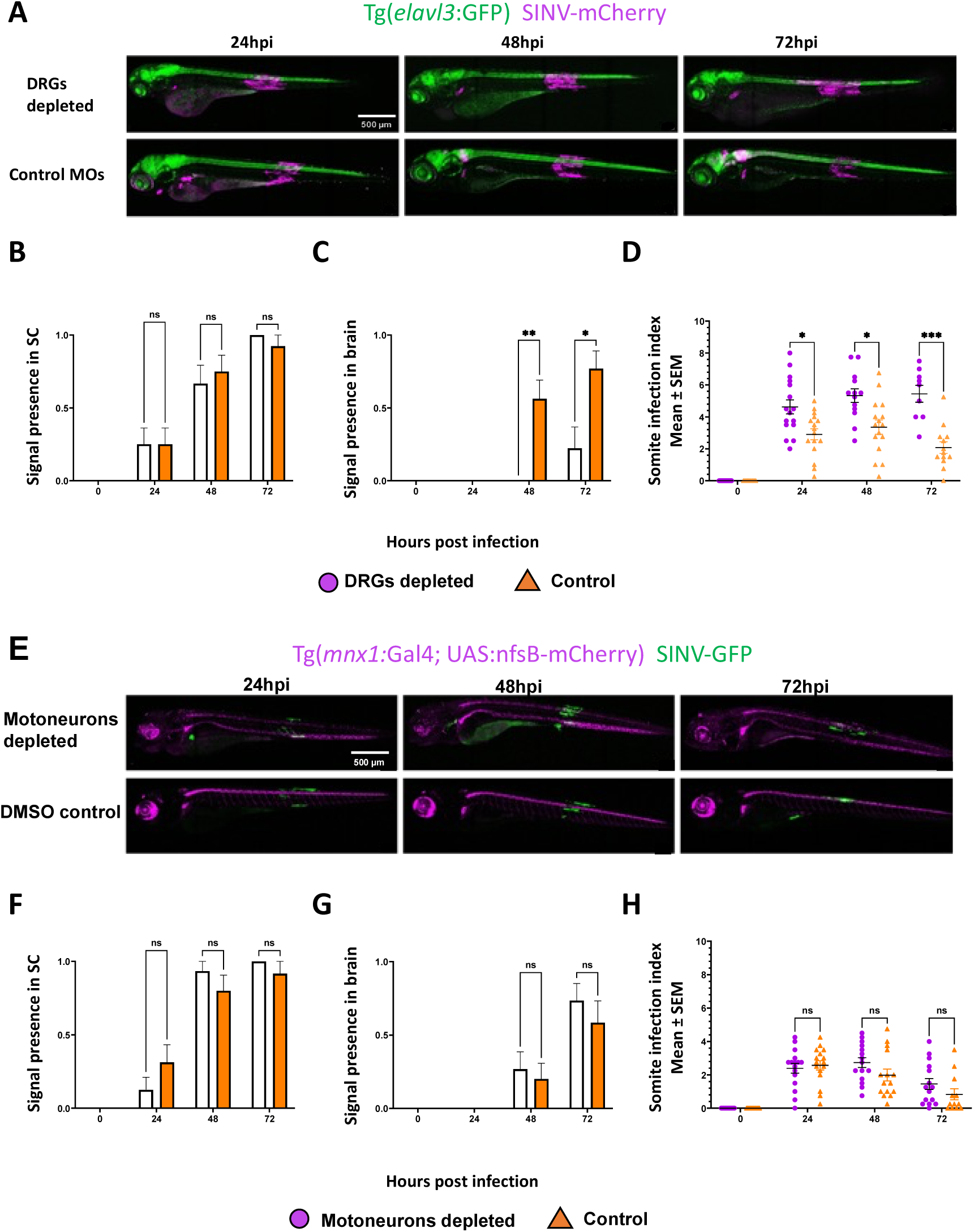
Impact of depletion of motor or sensory neurons on SINV infection. **A)** Maximum projection of confocal images of Tg(evlav3:GFP) (green) infected IM with SINV-mCherry (magenta) with DRG depleted by erbb3b morpholino treatment. **B-C)** Frequencies of spinal cord and of brain infection and **C)** Quantification of somite infection in DRG-depleted larvae followed over 3 days (n=16, 2 independent experiments) **D)** Maximum projection of confocal images of Tg(mnx1:Gal4, UAS:nfsB-mCherry) IM with SINV-GFP with motoneurons depleted by metronidazole treatment. **F-G)** Frequencies of spinal cord and of brain infection and **H)** Quantification of somite infection in motoneuron-depleted larvae followed over 3 days (n=16, 2 independent experiments). (***P < 0.001; **P < 0.01; *P < 0.05; ns - not significant)

Unexpectedly, somite infection was more persistent in DRG-depleted, but not motoneuron-depleted larvae (Figure 4D,H).

### SINV induces a strong IFN response, associated with transient peripheral infection

After having analyzed the dissemination patterns of the virus by real time microscopy, we performed quantitative measurements of infection dynamics after IM inoculation.

First, we performed qRT-PCR on whole larval lysates to measure the expression of viral and host genes. Strong expression of viral transcripts was measured at 24 hpi, remained stable at 48 hpi; a stronger dispersion was observed at 72 hpi, with a >10-fold decrease in most animals but no decrease in a few others (Figure 5A). A strong type I IFN response was induced, with a sustained *ifnphi1* induction (Figure 5B) but transient *ifnphi3* induction (Figure 5C). Accordingly, *MXA*, which is an IFN-stimulated-gene (ISG) inducible by both IFNф1 and IFNф3 (Aggad et al. 2009), was induced in a strong, sustained manner (Figure 5D).

**Figure 5:**
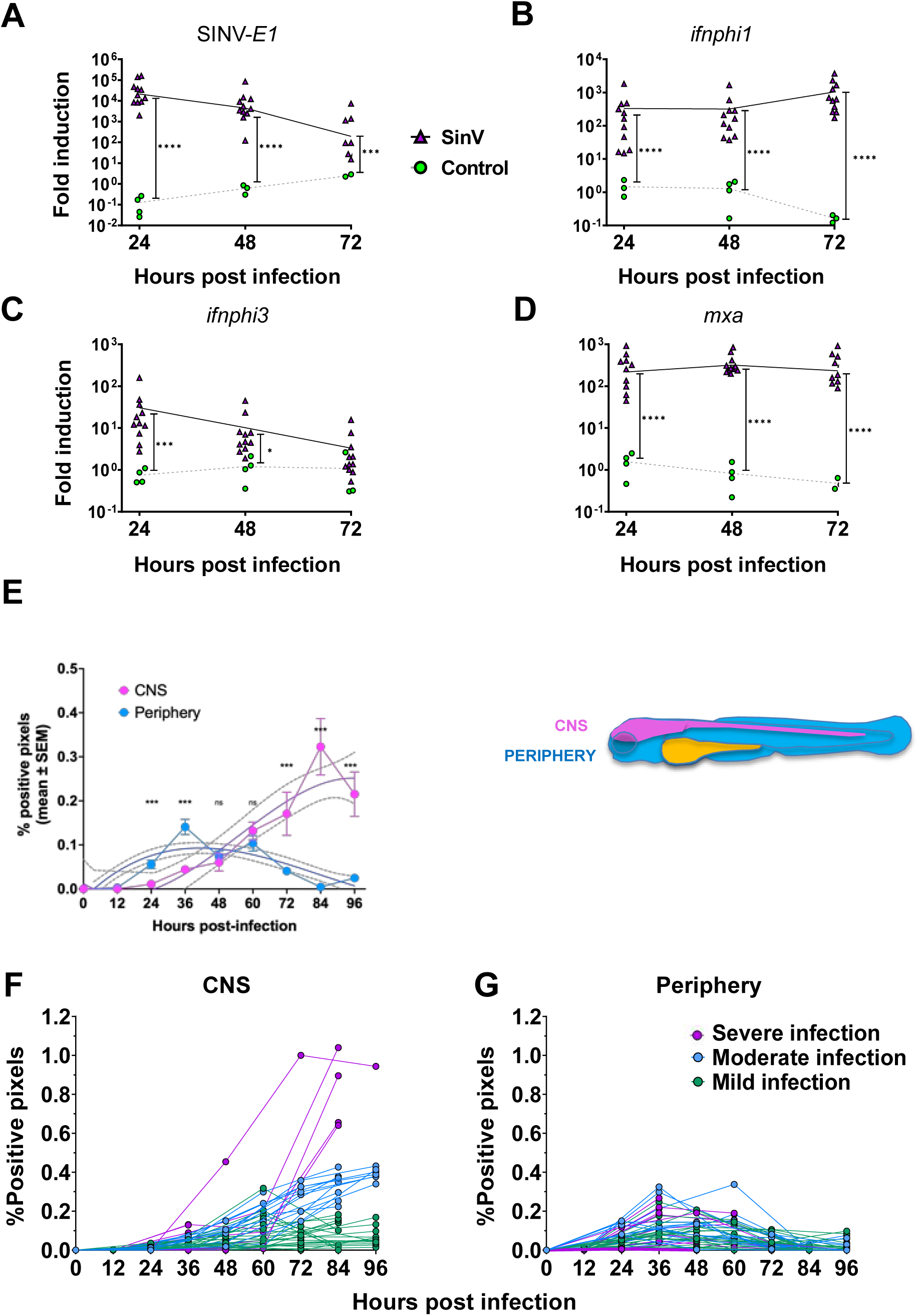
Kinetics of SINV infection and host response in zebrafish larvae. **A-D)** Real time quantification of genes expression of SINV-E1, infphi1, ifnphi3 and mxa of WT zebrafish larvae infected with SINV-GFP or SINV-mCherry IM and sampled every 24 hours (n=12, 2 independent experiments pooled) **E-G)** Pixel quantification of SINV-GFP signal percentage in CNS (Magenta) or periphery (Cyan) in WT larvae imaged every 24 hours (n=48, 4 independent experiments). E: collated values with polynomial interpolation and 95% confidence interval; F and G: individual trajectories (***P < 0.001; **P < 0.01; *P < 0.05; not shown - not significant)

After this whole-body analysis, to distinguish the dynamics of infection in periphery from CNS, we designed a semi-automatic 2D image analysis pipeline (detailed in methods). By segmenting the virus-encoded fluorescence signal from areas corresponding to CNS (spinal cord and brain) and periphery (rest of the body, excluding yolk and eyes), we quantified the extent of infection in these two compartments over time (Figure 5E-G). We verified that the data generated by this image analysis pipeline fitted with qPCR-based quantification by infecting larvae for one to three days and imaging them before lysis and RNA extraction; the sum of periphery+CNS positive pixels was remarkably well correlated with *SINV-E1* transcripts quantification in the whole larva (Figure S3).

As expected from our initial observations (Figure 1), infection in periphery was quantitatively transient, with a peak of fluorescence around 36 hpi, while infection in the CNS started later and increased until 4 dpi, the latest point analyzed (Figure 5E). Interestingly, the variance of the signal in the periphery was relatively low during the whole time-course, while it greatly increased with time in the CNS. This reflects a strong heterogeneity in individual courses of infection of the CNS, with some fish that appeared to control the infection while it soared in other ones (Figure 5F). Comparatively, infection course in periphery was quite homogeneous (Figure 5G).

In conclusion, infection in the periphery is transient and consistent among individuals, with a peak between 36 and 48 hpi. By contrast, CNS infection is progressive and more variable.

### Type I IFN response is systemic in periphery but mainly restricted to myeloid cells in CNS

To better understand this difference between peripheral and CNS compartments, we followed over time by live microscopy the IFN induction using a IFN response reporter line infected with SINV. Tg(*MXA*:mCherry) larvae inoculated with SINV-GFP displayed strongly elevated overall mCherry fluorescence (Figure 6A,B), with a 24h-delay relative to endogenous *MXA* expression (Figure 5D) as previously observed with other reporter transgenes (Palha et al. 2013). Virtual sectioning (Figure 6C) revealed that the *MXA*:mCherry transgene was highly induced in all gut, liver and skin – even in areas very distant from the infection, such as the jaw. However, in the spinal cord, only a few positive cells are detected, always in the vicinity of infected neurons but distinct from them. Therefore, response to IFN is systemic in peripheral epithelia but seems to be restricted to a few cells in the spinal cord. Similarly, following the spatial dynamics of the IFN response using the Tg(*ifnphi1*:mCherry) reporter fish revealed very few positive cells inside the spinal cord, despite an increase of mCherry positive leukocytes at the site of peripheral infection and, to a lesser extent in the entire body (Figure S4). Time-lapse imaging of scattered MXA or ifnphi1-reporter positive cells in the periphery or in the spinal cord revealed that they were mobile, suggesting they could be leukocytes (not shown). Thus, we imaged macrophage reporter fish Tg(*mfap4*:mCherry). While we could not detect microglia or macrophages inside the spinal cord of uninfected larvae, a few mCherry positive cells were observed in proximity to infected spinal cord areas. Using time-lapse, we could image macrophages entering the infected spinal cord, with directionality towards the most infected areas (Movie S10, Figure 6D).

**Figure 6.**
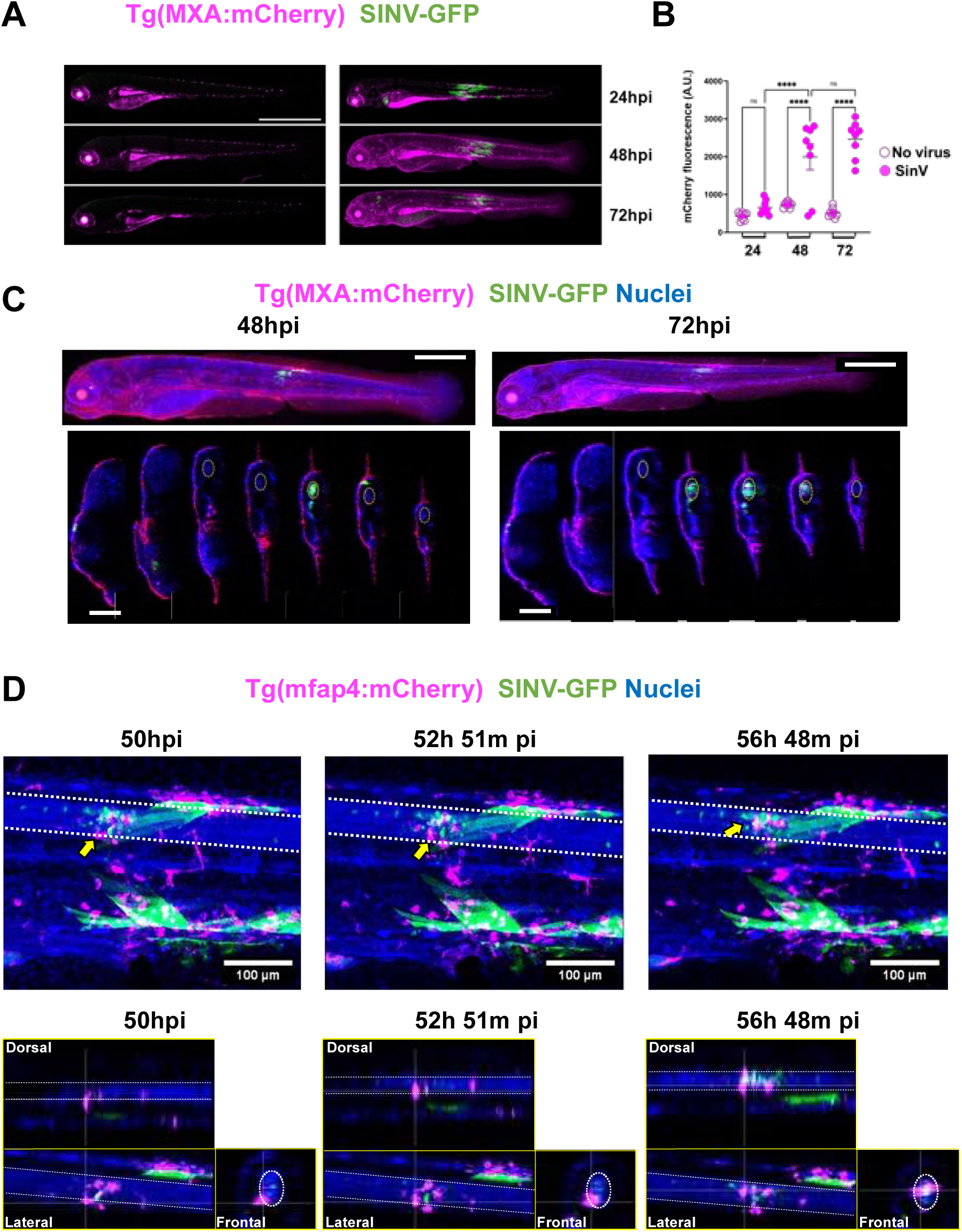
Spatial features of IFN response during SINV infection. **A-C)** Tg(MXA:mCherry) (magenta) infected IM with SINV-GFP (green). **A**. Maximum projection of confocal images of representative individual larvae over time. Scale bar, 1mm **B)** Mean intensity measurement of mCherry signal over the whole larva (n=8 per group). **C)** Maximum projection of confocal images of a larva counterstained by NucRed live (blue) (top) and relative transverse sections (bottom). Spinal cord indicated by the dotted oval on sections. Scale bar, 500µm on top, 100µm on bottom. **D)** Maximum projection of confocal images of Tg(mfap4:mCherry) (magenta), infected IM with SINV-GFP (green), nuclei in blue, (top) and relative orthogonal projections (bottom). Spinal cord limits indicated by dotted lines. The yellow arrow points to a macrophage entering the spinal cord. (***P < 0.001; **P < 0.01; *P < 0.05; ns - not significant)

In conclusion, while we observe a systemic type I response in the periphery, only very few IFN or ISG-expressing cells were observed in the spinal cord in the vicinity of highly infected areas, and our data strongly suggest that these cells are macrophages attracted to the infection site.

### Type I IFN responses are essential in controlling SINV infection

To delineate the role of this type I IFN response, we first used morpholinos to knock-down type I IFN receptors, previously shown to be highly efficient (Levraud et al. 2019; Palha et al. 2013). IFNR morphant larvae IM-injected with SINV displayed a more widespread and severe infection (Figure 7A) with strong mortality (Figure 7B). Accordingly, qRT-PCR analysis revealed a higher viral burden (Figure 7C). IFN induction was stronger in IFNR morphants, as predicted from the higher viral burden (Figure 7D,E). MXA induction was lower at 24 hours (Figure 6F), as expected with IFN-R knockdown; at 48 and 72 hpi, it was similar to that of controls, still consistent with IFN-R knockdown when considering stronger type I IFN expression.

**Figure 7:**
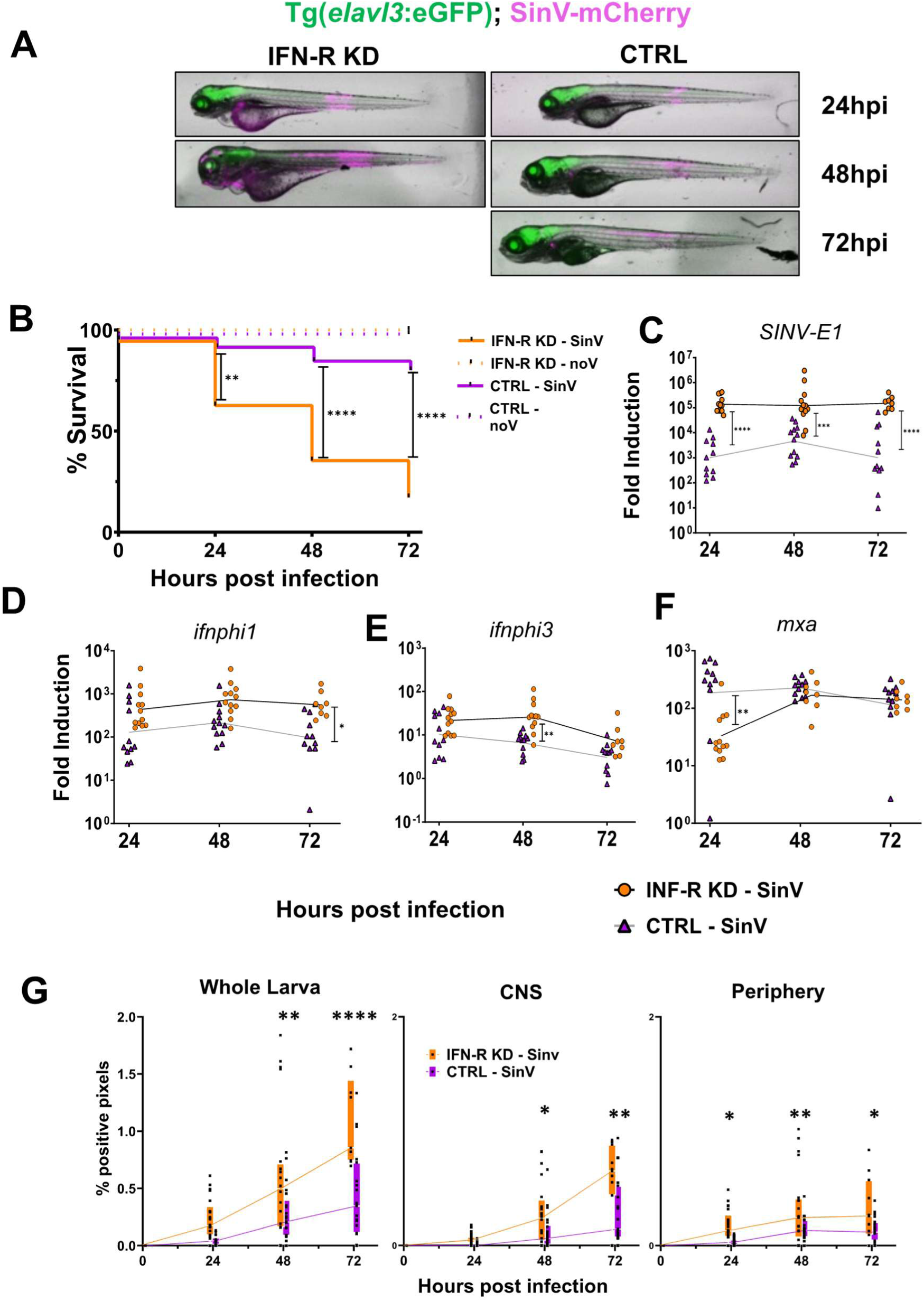
SINV kinetics and host gene expression of IFNR knock-down larvae. ***A)*** Top: Significative fluorescent images of Tg(elavl3:GFP) larvae infected with 30 PFU SinV-mCherry IM and injected either with IFNR or Control morpholino. The same larvae were followed each day. Bottom: Survival curve of Interferon Knock-down, wild type infected and not infected fish with 30 PFU SinV-mCherry IM. (n=44, 2 independent experiments) **B)** Real time quantification of gene expressions of SINV-E1, infphi1, infphi3 and mxa of zebrafish larvae infected with 30 PFU SINV-mCherry IM, or uninfected (control) injected with either IFNR or control morpholinos and sampled every 24 hours from the same experimental pool. (n=12, 2 independent experiments) **C)** Pixel quantification of SINV-mCherry signal percentage in whole body (Left), CNS (Center) or periphery (Right) of IFN KD and WT larvae. Injection of 30 PFU IM (n=21, 2 independent exp) (***P < 0.001; **P < 0.01; *P < 0.05; not shown - not significant).

To quantify the impact of IFNR knockdown in CNS and periphery, we used the image analysis pipeline introduced earlier. Infection was quantitatively stronger both in CNS and in peripheral compartments. In IFNR morphants, infection stabilizes but does not decrease in the periphery, while massive CNS infection systematically occurs (Figure 7G).

In our previous analysis of route of SINV entry to the CNS (Passoni et al. 2017), we had observed that endothelial cells were not infected by SINV, excluding CNS entry via BBB infection. We verified if endothelium infection occurred in IFNR morphants using Tg(*Fli1*:GFP) larvae infected with SINV-mCherry. After fixation and tissue clearing, high-resolution imaging did not reveal any infection of endothelial cells, even in strongly infected IFNR morphants (Movie S11).

To confirm the results obtained with morpholinos, we inoculated SINV-mCherry IM to “triple-phi” mutants carrying null mutations in *ifnphi1*, *2* and *3* genes (Laghi et al. 2022). As expected, triple-phi mutants were highly permissive for SINV propagation. Imaging, survival, and qRT-PCR analysis of the response were entirely consistent with those found in IFN-R morphants (Figure S5 and S6). Viral burden was higher in the mutants both in periphery and in the CNS (Figure S5D).

Together, these observations demonstrate that type I IFN responses play an essential role in controlling SINV infection, both in periphery and in the CNS.

### Mathematical modeling of the infection

To get insights into the in vivo dynamics of the infection in the whole animal, we generated a differential equation-based model, based on (Best et al. 2017), as detailed in Supplementary Text 2. The model considers two largely separate compartments, periphery and CNS. Briefly, each compartment includes three cellular subsets (Figure 8A): uninfected target cells, infected cells in eclipse phase (still functional and not yet producing virus), and productively infected cells entirely diverted into making new virions. Each compartment also has a pool of free virus and a pool of type I IFN. IFN induction pathways are highly conserved in zebrafish and mammals (Langevin et al. 2013). We considered two ways that IFN may be produced: via the Rig-I-like receptor (RLR) pathway, detecting of viral RNA in the cell cytosol of eclipse phase cells, or via the Toll-like receptor (TLR) pathway, detecting free virions. We also considered three ways that IFN may counteract the infection: by preventing virus from successfully entering a target, by preventing eclipse phase cells from entering productive state, and by accelerating the death of productively infected cells. Ignoring the effect of IFN amounted to modeling infection in IFNR-deficient larvae.

**Figure 8.**
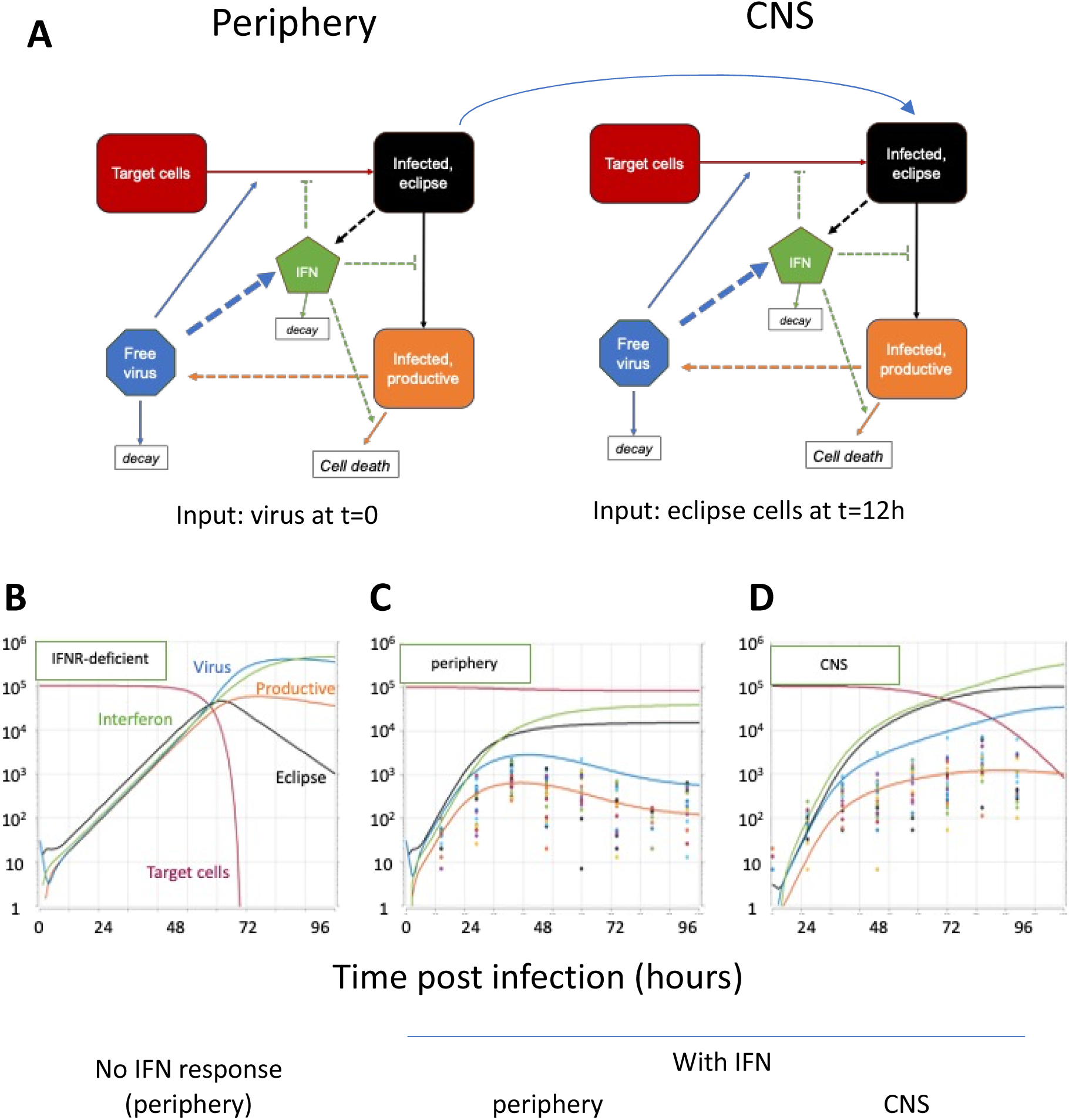
Modelling SINV infection in zebrafish larvae (see annex for details). A. organization of the model: periphery and CNS are ran as independent compartments, each with five variables: number of uninfected target cells, of infected cells in eclipse phase, and of productively infected cells; quantity of free infective virions; and concentration of type I IFNs. B-D results of running the model after parameter fitting. Initial conditions: all variables set to zero except virus=30 at t=0 for periphery, and eclipse cells=3 at t=0 (corresponding to 12hpi) for CNS. B. Peripheral compartment if IFN has no effect (ie as in IFNR morphants). C. Peripheral compartment in wt larvae; D. CNS compartment in wt larvae. Open circles in C and D derive from individual mage-based measure of infection displayed on Figure 5 F/G (from 48 larvae).

We fitted the parameters based on available literature and our own experimental data. To better translate our image quantification data into amount of infected cells, we performed flow cytometry (FC) to estimate the fraction of infected cells in larvae. Fluorescent reporters in our SINV strains are co-expressed with viral structural proteins, which are expressed during the late, productive stage of the infection (Strauss and Strauss 1994a). Control and IFNR-morphant larvae were infected IM with SINV-GFP for one to three days, imaged, then dissociated to assess the fraction of GFP positive cells by FC. Imaging and FC results were well correlated (Figure 8 supplement 1), although not as well as imaging and qRT-PCR data (Figure 5 supplement 1), possibly because dissociation efficiency is tissue-dependent, introducing biases. This, however, revealed that in the strongest infections, up to 20% of the cells may become GFP positive. Since not all larval cells are infectable by the virus (e.g, endothelial cells) we estimated that death would occur when half of target cells had become productively infected in the periphery.

One parameter that was difficult to assess was the rate of new virus production per infected cell. However, plausible values could be attributed to other parameters unrelated to IFN response. In IFNR-deficient larvae, rate of virus production determines when this threshold of 50% infected cells is reached. Considering that IFNR-deficient larvae typically die in two to three days from the infection, we could estimate the rate of new virus release by productively infected cells, which was surprisingly low: approximately 1 virus per hour and per cell. The corresponding simulation is displayed on Figure 8B.

To fit the parameters of IFN response in periphery, we used the fact that in IFN-competent fish, infection peaks between 24 and 48 hpi (Figure 5G and S5D); our FC data suggest that at this time point, approximately 1% of the cells should be productively infected. We found parameter values that reasonably recapitulated this peak and the following decay. For this decay to occur, the RLR pathway was required, and IFN had to accelerate the death of infected cells (see annex). We then refined the values by performing multi-parameter fitting on the actual infection data of the periphery of 48 larvae; the resulting curves are shown on Figure 8C, and were fully consistent with our previous assumptions.

These parameters result in a robust control of the infection, which is the situation observed in periphery. The situation in the CNS was different and parameters had to be adjusted to better reflect our results. We assumed that the BBB prevents the passage of IFN or free virions between the two compartments; the same overall model is therefore used for the CNS. Since we detected the first SINV-fluorescent neurons around or before 24 hpi, and determined statistically an average of 3 independent CNS entry events, the CNS simulation starts with 3 neurons entering eclipse phase at 12 hpi.

Since death of infected neurons was rare, we expected to find a smaller value for CPE-induced death in CNS. Also, based on known properties of neurons (Viengkhou and Hofer 2023) and our results with MXA and IFN reporter fish, we expected a less efficient RLR pathway, and an overall smaller impact of the IFN response on infected neurons. We performed multiparameter fitting on the CNS infection values previously measured (Figure 8D). Remarkably, in this best fit solution, IFN is mostly produced via the TLR pathway, and IFN main impact is to prevent exit from eclipse phase, which is predicted to be the fate of almost CNS cells. This would be as close as possible to the variable outcomes we observed, with CNS infection being sometimes controlled and sometimes not, in a model that is by essence deterministic.

In conclusion, despite its limitations, our model unveiled plausible values for two unknown parameters of SINV infection in zebrafish: productively infected cells release approximately only one infectious virion per hour, and the IFN response is ten times less efficient in the CNS than it is in periphery.

## Discussion

Taking advantage of the exquisite imaging assets of zebrafish larvae, we describe here in detail the spatial dynamics of a viral infection as it propagates from periphery to CNS in a whole vertebrate. Our previous work had established that SINV is neuroinvasive in zebrafish (Passoni et al. 2017). This had been performed using IV injection, yielding highly variable patterns of infection. Here, we compared several inoculation sites, and found that this variability can be reduced; interestingly, CNS invasion always occurred. When the virus was injected in the pericardial cavity, the infection was confined to the anterior half, with invasion of the hindbrain, probably via cranial ganglia. Upon IM injection in a caudal somite, peripheral infection remained localized to the tail, from where the virus invaded the spinal cord, and sometimes later spread to the brain. Subsequently, we focused on the outcome of IM injection, which reflected most closely natural SINV transmission by a mosquito bite. Furthermore, the thinness of the tail favored high-resolution imaging, and the relatively simple organization of the spinal cord facilitated the analysis of CNS entry and propagation. After IM injection, SINV showed a strong tropism for muscle fibers and for stromal cells between somites or close to ventral or dorsal vessels, but not for endothelial cells themselves. This muscular tropism is relevant to the clinical features of SINV infection in humans, who sometimes develop long-lasting myalgia, associated with cycles of necrosis and regeneration in the patient muscles (Sane et al. 2012). Indeed, we observed rapid death of infected cells in zebrafish muscle, but not in a cyclical manner; in the 4-day time frame of our observations, peripheral infection appeared to be fully resolved. Although we did not analyze it in detail, muscle regeneration clearly occurred, not surprisingly considering the naturally potent regeneration abilities of zebrafish larvae. However, when IFN receptors were knocked down, the larvae lost the ability to control virus propagation in muscle, resulting in widespread areas of infection over multiple somites and subsequent extensive necrosis.

Spinal cord invasion always starts in the segment that innervates the infected somite, which is entirely consistent with CNS entry by axonal transport, as demonstrated previously (Passoni et al. 2017). As this could involve either motor or sensory neurons, we used time lapse imaging to determine which became infected first. We observed both situations – and since we have also established that, on average, 3 independent CNS entries occurred per larva, entry may take place by both sensory and motor neurons in the same larva. To establish the relative importance of both gateways, we performed depletion experiments. Spinal cord invasion still occurred after elimination of either sensory or motor neurons, confirming that both populations may constitute the entry route to the CNS. Still, some remarkable differences were observed after sensory or motor neuron depletion. Most strikingly, subsequent spread of SINV to the brain was suppressed when DRG sensory neurons were depleted. This would be expected if long-distance axonal propagation of SINV was more efficient by anterograde than by retrograde transport. Surprisingly, DRG neuron depletion also had an impact on the periphery, resulting in more extensive somite infection. This suggests that DRG sensory neurons may somehow contribute to the antiviral response, possibly by playing an important immuno-sensing role. However, DRG depletion was achieved by knocking down *erbb3b*, a transcription factor expressed in neural crest progenitors, preventing them to differentiate into sensory neurons or melanophores (Dooley et al. 2013), raising the possibility that another unknown neural crest-derived cell type may prevent virus spread.

High-resolution imaging clearly showed that most, if not all, cells infected in the spinal cord were neurons; the other prominent cell type, radial glia, would have been readily identified by their typical morphology if they had become fluorescent upon infection. We observed two modalities of virus dissemination inside the CNS: axon-mediated long-distance transport (almost always in the anterior direction) and local cell-to-cell spread, most likely by virus shedding from neuronal soma. This resulted in the formation of progressively growing clusters. Cluster formation was particularly evident in larvae directly inoculated in the spinal cord with a mixture of SINV-GFP and SINV-mCherry, where stable territories of green and red fluorescent neurons became established without mixing. This remarkable virus exclusion phenomenon was observed everywhere, with the notable exception of the yolk. Superinfection exclusion has been well documented for SINV, requiring at most 1h after initial infection to be established (Strauss and Strauss 1994b). Unlike other cells, the very large size of the yolk syncytial cell makes it likely that it could be infected by several virions simultaneously, and more time would also be required for exclusion signals to travel to the entire cell.

This cluster growth played a clear role in progression of infection in the CNS, but axonal long-distance spread played a decisive role, seeding new distant clusters in the spinal cord and, more importantly, in the brain, starting with the hindbrain where the targets of spinal neurons reside. CNS infection was highly variable among individuals, with some larvae that managed to control the virus and others where a runaway infection occurred; apparition of new clusters in the anterior spine and in the brain was a major difference between the two groups. What is causing this initial difference is unclear but clearly of critical importance; as stated above, initial entry via sensory neurons, rather than motoneurons, is likely to be a major determinant. SINV induces a strong type I IFN response in larvae. The dynamics of this response, as revealed by qRT-PCR in the whole larvae after IM inoculation, are very similar to those measured after IV infection of SINV (Boucontet et al. 2018) or of its relative CHIKV (Palha et al. 2013). The effect of knocking down IFN receptors was also similar, with a much stronger overall infection resulting in the death of most larvae within ∼3 days. Similar results were obtained when stable mutants lacking the type I IFN genes ifnphi1, 2 and 3 (triple phi) were infected. We used reporter transgenes to assess the localization of IFN responses. ISG expression, as determined in Tg(MXA:mCherry) larvae, was highest in the gut and the skin; it was also high in leukocytes, particularly near to infected sites. Skin expression was remarkably uniform, being detected even in regions farthest from the infection, revealing a systemic IFN response in the periphery. This probably explains why SINV infection is efficiently controlled in the periphery, with a stereotypic transient infection. Interestingly, in IV-inoculated larvae, we observed at least one exception to this: when infection occurred in the swim bladder, it was highly persistent, which would be worth of future investigation. The situation in the CNS was very different, with MXA reporter expressing being restricted to a few mobile cells found only in already extensive clusters of infected neurons. It is very likely that these cells are macrophages attracted from the outside. Indeed, we could document the entrance of macrophages into the spinal cord, with a clear directional tropism towards the infected clusters. What is attracting these macrophages would be an important question to address in the future.

To get deeper insights into the mechanisms by which the IFN response counteracts the infection in the periphery and in the CNS, we resorted to mathematical modeling. The parameters of our differential equation model were first determined for the peripheral compartment. After establishing various parameters based on literature or our literature data, we used simulations to fit the remaining ones. To match the survival time of IFN-R knockdown larvae, our model yielded an unexpected result for the rate of virus release by productively infected cells: about one infective virion per hour. This may appear very low since transmission electron microscopy of SINV-infected cells typically reveals many virions budding simultaneously at the plasma membrane; time-lapse imaging of tagged viruses detected several budding events per minute (Jose et al. 2015). However, it should be kept in mind that not all viral particles are infective virions; actually, comparisons of PFU titers and amount of viral genomes in typical BHK-produced SINV suspensions indicate than less than 1% of particles are infective (Poirier et al. 2015). Thus, this rate of 1 virion/infected cell/hour is not unrealistic; and it is sufficient to yield an explosive increase of the infection, if not counteracted by host immunity.

Our model allowed for various modalities of IFN production and impact of IFN on the infection. Testing these in turn did not allow to reach definitive conclusions, but indicated that the RLR detection pathway was important, and that IFN is probably accelerating the death of infected cells, as we hypothesized previously (Levraud et al. 2014). One notable difference in our simulations, when comparing the different possible impacts of IFN, was in the number of cells in eclipse phase (see Annex). These are currently undetectable with our imaging tools and developing means to visualize them would be important to test key hypotheses regarding infection dynamics in vivo.

The model also suggested that IFN were much less efficient on neurons than on peripheral cells. As far as we know this has not been compared directly, but it is well established that non-neuronal cells of the CNS (e.g. microglia or astrocytes) have a stronger response to IFN than neurons; furthermore, the repertoire of ISGs produced by neurons is relatively narrow (Viengkhou and Hofer 2023). This can be understood as a tradeoff between the need to protect the CNS from viral infection and the deleterious impacts of neuroinflammation.

Our mathematical modeling has important limitations. First, our choice of parameters is sometimes arbitrary, if plausible, and will have to be refined in the future. The equations governing eclipse cells are clearly too simple, since in presence of a strong IFN response, our model induce eclipse cells to remain in this state forever; a return pathway to the target state, or to a transient refractory state, should be included in a more realistic – but more complex – model. Most important for the CNS component of our model, the localization IFN response is not taken in account. In all likelihood, local neuron-to-neuron spread of the virus is soon counteracted by the local IFN response; but when axonal transfer result in a new infection focus at a very distant site, this new area is probably completely unprotected for some time. It would be relevant to adapt this approach to drug discovery, possibly with the help of artificial intelligence to better discover homologies between effective patterns and predicted drug effects.

To conclude, this study analyzing the dynamics of viral spreading and IFN responses in real time proved of high value to guide modelling and reveal novel features governing dissemination of viruses through the whole organisms, and particularly for the very important events of CNS invasion.

## Material and methods

### Ethical Statement

Animal experiments described in the present study were conducted according to European Union guidelines for handling of laboratory animals and were approved by the Ethics Committees of Institut Pasteur and Paris-Saclay Neuroscience Institute.

### Zebrafish lines

Zebrafish were handled as explained in (Laghi et al., 2022). Wild-type zebrafish (AB strain), initially obtained from ZIRC (Eugene, OR, USA), were raised in the aquatic facility of Institut Pasteur. After natural spawning, eggs were collected, treated for 5 min with 0.03% bleach, rinsed twice, and incubated at 28°C in Petri dishes in Volvic mineral water supplemented with 0.3 µg/ml methylene blue (Sigma-Aldrich, St. Louis, Missouri, USA). After 24 h, the water was supplemented with 200 µM phenylthiourea (PTU, Sigma-Aldrich) to prevent pigmentation of larvae. After this step, incubation was conducted at 24°C or 28°C depending on the desired developmental speed. Developmental stages given in the text correspond to the 28.5°C reference (C B Kimmel et al., 1995). At 3 dpf, immediately before infections, larvae that had not hatched spontaneously were manually dechorionated. Owing to silencing issues of some UAS-driven transgenes, breeders were carefully screened to select those whose progeny yielded full expression; correct fluorescence expression by larvae was checked before experiments. Sex of larvae is not yet determined at the time of experiments. A list of the transgenic zebrafish lines used in this experiment and their reference can be find in Table 1.

**Table 1.** Transgenic zebrafish.

| Organism | Line Name | Modification | Transgene name and reference | Reference |
|---|---|---|---|---|
| *Danio rerio* | Mnx1::Gal4 | Transgenesis | **Tg1(mnx1:GAL4)** https://zfin.org/ZDB-TGCONSTRCT-160119-3 | https://zfin.org/ZDB-TGCONSTRCT-160119-3 |
| *Danio rerio* | axl1::eGFP | Transgenesis | **Tg(foxa2:GFP)** https://zfin.org/ZDB-GENE-980526-404 | https://zfin.org/ZDB-GENE-980526-404 |
| *Danio rerio* | IFN1::mCherry allele | Transgenesis | **Tg(ifnphi1:mCherry)** https://zfin.org/ZDB-TGCONSTRCT-120607-1 | https://zfin.org/ZDB-TGCONSTRCT-120607-1 |
| *Danio rerio* | HuC-GFP | Transgenesis | **Tg(elavl3:EGFP)** https://zfin.org/ZDB-TGCONSTRCT-070117-150 | https://zfin.org/ZDB-TGCONSTRCT-070117-150 |
| *Danio rerio* | Huc:Gal4 | Transgenesis | **Tg6(elavl3:GAL4-VP16)** https://zfin.org/ZDB-ALT-161116-3 | https://zfin.org/ZDB-ALT-161116-3 |
| *Danio rerio* | Ml2Tg (hb9::eGFP) | Transgenesis | **Tg(mnx1:GFP)** https://zfin.org/ZDB-TGCONSTRCT-070117-40 | ttps://zfin.org/ZDB-TGCONSTRCT-070117-40 |
| *Danio rerio* | Sb1Tg (ngn1::eGFP) | Transgenesis | **Tg(ngn1::eGFP)** https://zfin.org/ZDB-GENE-990415-174 | https://zfin.org/ZDB-GENE-990415-174 |
| *Danio rerio* | MXA:mCherrry | Transgenesis | Tg(cryaa:DsRed,mxa:mCherryF) | http://zfin.org/ZDB-ALT-190116-3 |
| *Danio rerio* | UAS:nfsB-mCherry | Transgenesis | UAS-E1B:NTR-mCherry c264; http://zfin.org/ZDB-FISH-150901-2592 | http://zfin.org/ZDB-FISH-150901-2592 |

### Viruses

Sindbis viruses were produced on BHK cells [originally obtained from American Type Culture Collection (ATCC), #CC-L10], according to (Hardwick and Levine, 2000). The SINV-GFP strain used here corresponds to the SINV-eGFP/2A strain described in (Boucontet et al., 2018), whose genome is based on the hybrid TE12 strain backbone. The SINV-mCherry strain was generated on the same backbone, replacing the eGFP coding sequence with mCherry. Viruses were titered on Vero-E6 cells (ATCC #CRL-1586).

### Injections

Injections and handling of larvae were performed as in (Passoni et al., 2017). Briefly, zebrafish larvae aged 70-72 hpf were inoculated by microinjection of ∼30 PFU viral SINV particles (∼1 nl of supernatant from infected BHK cells, diluted with PBS to ∼3x10^7^ PFU/ml). Before injection larvae were anesthetized with 0.2 mg/ml tricaine and positioned and oriented in the groove molded in 2% low melting agarose. Using a micromanipulator, the capillary was then inserted at the desired site and two pulses were performed to inject approximately 1 nl. The larvae were then distributed in wells of culture plates with water containing PTU and kept at 28°C. For ethical reasons, all larvae used in the experiments were euthanized by anesthetic overdose at 7 dpi.

### Lysis, RNA Extraction, and RT-qPCR of Larvae

RNA was extracted from individual larvae, which were first deeply anesthetized. After the removal of almost all water and the addition of RLT buffer (Qiagen), larvae were dissociated by 5 up- and-down-pipetting movements. Tubes may then be frozen at -20°C for a few days. Total RNA was then extracted with a RNeasy Mini Kit (Qiagen) without the DNase treatment step and a final elution with 30 µl of water.

RT was performed on 6 µl of eluted RNA using MMLV reverse transcriptase (Invitrogen, Carlsbad, CA, USA) with dT_17_ primer (for polyadenylated transcripts). cDNA was diluted with water to a final volume of 100 µl, of which 5 µl was used as a template for each qPCR assay.

Real-time qPCR was performed with an ABI7300 (Applied Biosystems, Foster City, CA, USA). Quantification was performed using a SYBR assay using the Takyon SYBR Blue MasterMix (Eurogentec, Seraing, Belgium) with primer pairs (Table 2). These primers typically span exon boundaries to avoid amplification of contaminating genomic DNA. The *eef1a1l1* (also known as *ef1a*) was used as a housekeeping gene for normalization.

**Table 2.**
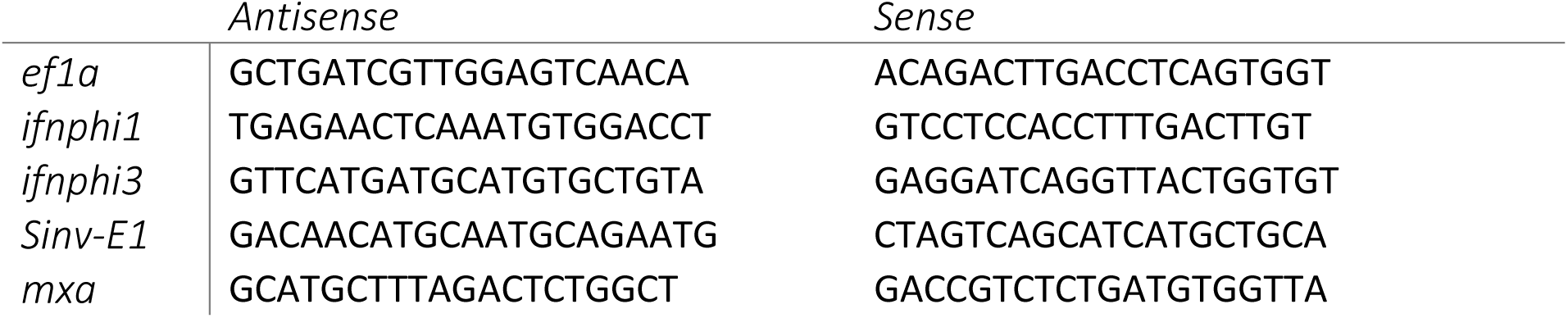
RT-qPCR primers.

### Morpholino and Plasmid Injection in Eggs

Morpholino antisense oligonucleotides (Gene Tools, Philomath, OR, USA) were injected (1 nl volume) in the cell or yolk of AB embryos at the one- to two-cell stage._crfb1 splice morpholino (2 ng, CGCCAAGATCATACCTGTAAAGTAA) was injected together with crfb2 splice morpholino (2 ng, CTATGAA TCCTCACCTAGGGTAAAC), knocking down all type I IFN receptors (Aggad et al., 2009). Erbb3b morpholino was used to prevent differentiation of DRG neurons and injected alone (2 ng, TGGGCTCGCAACTGGGTGGAAACAA) (Dooley et al., 2013). Control morphants were injected with 4 ng control morpholino, with no known target (GAAAGCATGGCATCTGGAT CATCGA).

### Live Widefield Fluorescence Imaging

Live fluorescence imaging was performed as explained in (Laghi et al., 2022). SINV-GFP-infected larvae were imaged with an EVOS FL Auto or EVOS M7000 microscope (Thermo Fisher Scientific, Waltham, MA, USA) using a 2× planachromatic objective (numerical aperture, 0.06), allowing capture of the entire larva in a field. Before imaging larvae were anesthetized with 0.2mg/ml tricaine and placed in individual wells of 24-well plate cell culture plate. Agarose 2% molds were used to properly set each larva in the same relative position. Transmitted light and fluorescence (GFP or Texas Red cube) images were taken. After imaging larvae were rinsed in water with 0.3 µg/ml methylene blue (Sigma-Aldrich, St. Louis, Missouri, USA) and transferred to individual wells of a 24-well plate cell culture plate in water with 200 µM phenylthiourea (PTU, Sigma-Aldrich) and stored in an incubator at 28 C.

### Live Confocal Imaging

Confocal imaging was performed as in (Viana et al., 2023). Injected larvae were mounted in lateral or ventral position in 35 mm glass-bottom-dishes (Ibidi Cat#: 81158) or in glass bottom-8well-slides (Ibidi Cat#: 80827). The larvae were immobilized using a 1% low-melting-point agarose (Promega; Cat#: V2111) solution and covered with Volvic water containing 0.2mg/ml tricaine. A Leica SP8 confocal microscope equipped with two PMT and Hybrid detector, a 10X dry (PL Fluotar 10X dry:0.30), 20X IMM (HC PL APO CS2 20X/0.75), or a 40x water IMM (HC PL APO CS2 40X/1.10) objective, an X–Y motorized stage and with LAS-X software, was used to live image injected larvae. To generate images of the whole larvae, a mosaic of confocal z-stack of images was taken with the 10X or 40X objective using the Navigator tool of the LAS-X software. The acquisitions were performed either in conventional settings or resonant scanning, the latter being post-processed with the Lightning tool of the LAS-X software to eliminate noise (deconvolution). After the acquisition, larvae were washed and transferred in a new 24-well plate filled with 1 ml of fresh Volvic water per well, incubated at 28°C, and imaged again under the same conditions over time.

Considering the image analysis strategy adopted and the need to adjust the acquisition setting to avoid over or under-saturation of the sensor, the strategy adopted to set acquisition parameters always followed the same rule. Every single channel was brought up to saturation of true signal (ignoring saturation of noise from yolk or eye) and subsequently lowered enough to remove the saturation. At every timepoint, this calibration is repeated on a batch of random samples and applied to the whole timepoint. In this way, the whole range of emission is captured for every image, without affecting the integrity of the subsequent threshold-based analysis.

### Image analysis

Image analysis was performed using FIJI software (Schindelin et al., 2012) and CellProfiler (Carpenter et al., 2006). Fluorescent images acquired through EVOS imaging were normalized (i.e., flip, crop, and denoising) using a FIJI script (Script 1). After normalization, a threshold check (biovoxxel toolbox plugin) was used to identify the best thresholding strategy for each channel, which resulted to be “Triangle dark” and “IsoData dark”. Subsequently, all the images were threshold using the same method (script 1). Both the binary masks and the normalized images were fed to CellProfiler pipeline (pipeline 1) to perform area filtering, segmentation, and measurement. All these operations were performed in batches. The final output obtained is a measurement of the positive pixel area normalized to the constant area of the image, segmented in CNS and Periphery. Only the tg(INFphi1:mCherry) experiment was quantified from confocal images and for this, all the operations were performed through CellProfiler software directly (pipeline2). In this case, the threshold was performed directly in CellProfiler using the “Otsu three class” method.

### Statistical analysis

The difference between means was evaluated using analysis of variance (ANOVA) or two-tailed unpaired t-test. For multiple comparisons, Bonferroni’s method was applied. For Normal distributions, the Kolmogorov-Smirnov test was used. Non-Gaussian data were analyzed with Kruskal-Walli’s test followed by Dunn’s Multiple comparison, except for qRT-PCR results, which were analyzed by lognormal ordinary ANOVA distribution to compare geometric means. P<0.05 was considered statistically significant (symbols: ***P<0.001; **P<0.01; *P<0.05). Survival data were plotted using the Kaplan-Meier estimator and log-rank tests were performed to assess differences between groups. Statistical analyses and PCA were performed using Prism software.

### Model building

Model building was realized using Berkeley Madonna software. This software numerically solves ordinary differential equations and difference equations. All the details on the parameters used in this software are accessible in the Annex.

## Acknowledgements

We thank Yohann Rolin for the zebrafish facility care, Valérie Briolat for help with protocols to analyze IFN responses, Benjamin tenOever and Ariel Benitez for expert advice on SINV inverse genetics, and Pierre Boudinot and Michaël Demarque for constructive discussions.

