## Supplementary figures, text, and movie legends for "Mapping the journey of a neuro-invasive virus in zebrafish larvae: contrasted viral spread and type I interferon responses in periphery and central nervous system"

Figure S1

Figure S2

Figure S3

Figure S4

Figure S5

Figure S6

Supplementary text 1

Supplementary text 2

Legends to supplementary movies

| | | JFG | SCA | FG | Br | Sp | n= | |
|---|---|---|---|---|---|---|---|---|
| d1 | green | 12 | 21 | 4 | 3 | 0 | 24 | INTRAVENOUS |
| | red | 7 | 21 | 1 | 0 | 0 | | |
| d2 | green | 13 | 20 | 10 | 11 | 5 | 23 | |
| | red | 11 | 18 | 5 | 9 | 7 | | |
| d3 | green | 10 | 16 | 8 | 15 | 9 | 22 | |
| | red | 7 | 17 | 3 | 11 | 14 | | |
| d4 | green | 0 | 7 | 4 | 16 | *9* | 22 | |
| | red | 2 | 12 | 3 | 13 | *17* | | |

| | | JFG | SCA | FG | Br | Sp | n= | |
|---|---|---|---|---|---|---|---|---|
| d1 | green | 24 | 0 | 2 | 2 | 0 | 24 | PERICARDIUM |
| | red | 21 | 0 | 1 | 0 | 0 | | |
| d2 | green | 24 | 0 | 3 | 6 | 0 | 24 | |
| | red | 24 | 0 | 4 | 7 | 0 | | |
| d3 | green | *8* | 0 | 2 | 11 | 0 | 24 | |
| | red | *18* | 0 | 4 | 11 | 0 | | |
| d4 | green | *6* | 0 | 1 | 13 | 1 | 23 | |
| | red | *20* | 0 | 4 | 13 | 0 | | |

| | | JFG | SCA | FG | Br | Sp | n= | |
|---|---|---|---|---|---|---|---|---|
| d1 | green | 4 | 23 | 1 | 0 | 0 | 24 | INTRAMUSCULAR |
| | red | 5 | 19 | 0 | 1 | 1 | | |
| d2 | green | 7 | 23 | 1 | 2 | 16 | 24 | |
| | red | 7 | 21 | 2 | 4 | 12 | | |
| d3 | green | 6 | 17 | 3 | 2 | 17 | 22 | |
| | red | 2 | 14 | 1 | 7 | 15 | | |
| d4 | green | 1 | 10 | 2 | 5 | 17 | 21 | |
| | red | 1 | 10 | 1 | 7 | 15 | | |

| | | JFG | SCA | FG | Br | Sp | n= | |
|---|---|---|---|---|---|---|---|---|
| d1 | green | 0 | 20 | 0 | 11 | 23 | 23 | SPINAL CORD |
| | red | 1 | 15 | 0 | 8 | 23 | | |
| d2 | green | 0 | 17 | 1 | 18 | 24 | 24 | |
| | red | 3 | 12 | 2 | 15 | 24 | | |
| d3 | green | 0 | 7 | 0 | 21 | 23 | 23 | |
| | red | 2 | 3 | 0 | 22 | 23 | | |
| d4 | green | 0 | 2 | 1 | 19 | 21 | 21 | |
| | red | 1 | 0 | 0 | 20 | 20 | | |

| | |
|---|---|
| x<br>y | blue background (plain)<br>no significant difference |
| *x*<br>*y* | yellow background (italics)<br>significant difference |
| Fisher's exact test | |

| | |
|---|---|
| **JFG** | Jaw, Gill, Facial muscles |
| **SCA** | Somites, caudal hematopoietic tissue, dorsal anastomotic zone |
| **FG** | facial ganglia |
| **Br** | brain |
| **Sp** | spinal cord |

**Figure S1.** Comparison of frequencies of infection by SINV-GFP and SINV-mCherry in five compartments, at different timepoints, in wild type larvae injected with a mix of 30 PFU SINV:GFP and SINV:mCherry by intravenous, intramuscular, pericardium or spinal cord inoculation. Differences in frequencies were considered to be statistically different for $p<0.05$ using Fisher's exact test.

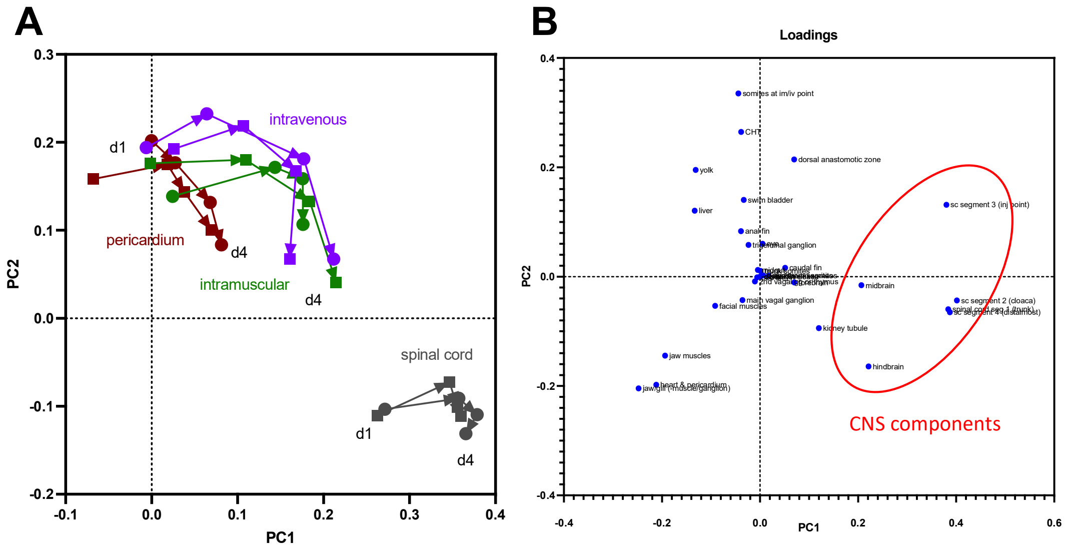

**Figure S2. A)** Principal component analysis of the frequency table of infection, comparing the different injection sites and the different time points (d1 and d4 correspond to day 1 and 4 post inoculation; arrows show time progression for a given group of larvae; intermediate days not shown for clarity). The two independently injected groups for each site were treated separately to assess reproducibility. **B)** table of weights (loadings) of each body region in the PCA analysis. The red oval contains CNS regions.

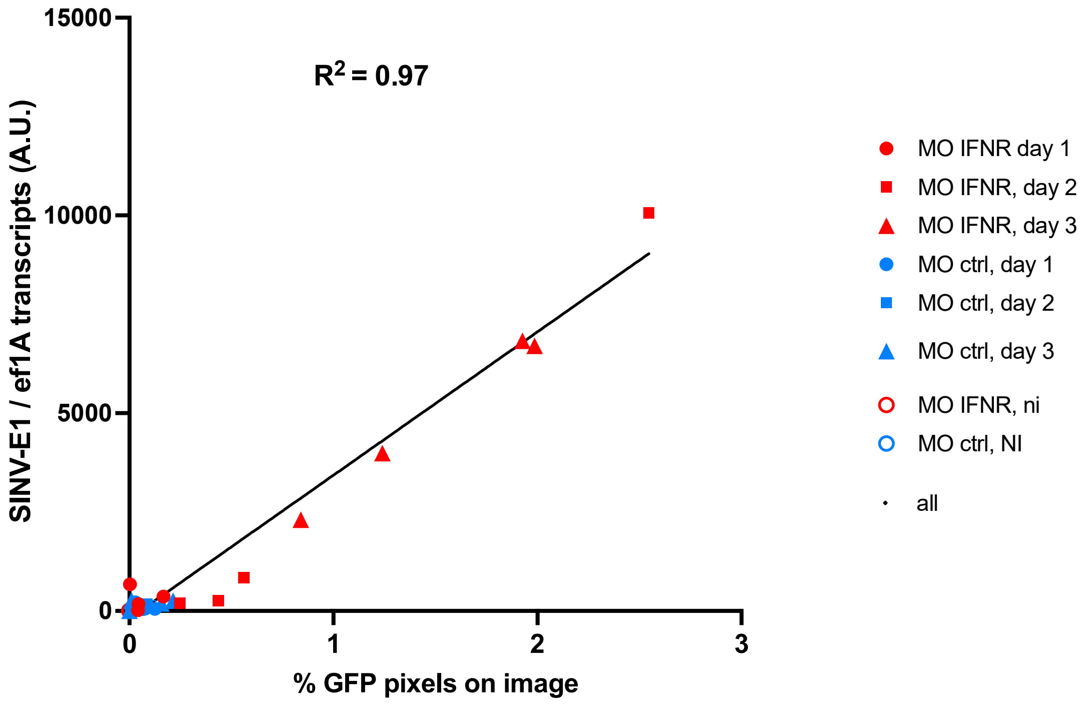

**Figure S3**. Correlation of viral burden measured by qRT-PCR and image quantification. Individual larvae, IFNR-proficient or deficient, were imaged at various times post-infection, and lysed immediately after for RNA extraction. Images were analyzed using the quantitative 2D analysis pipeline, and the fraction of positive pixels in CNS and periphery added to yield the x-value; while RT-qPCR quantification on viral E1 transcripts was performed to yield the y-value. N=23 infected fish + 6 uninfected.

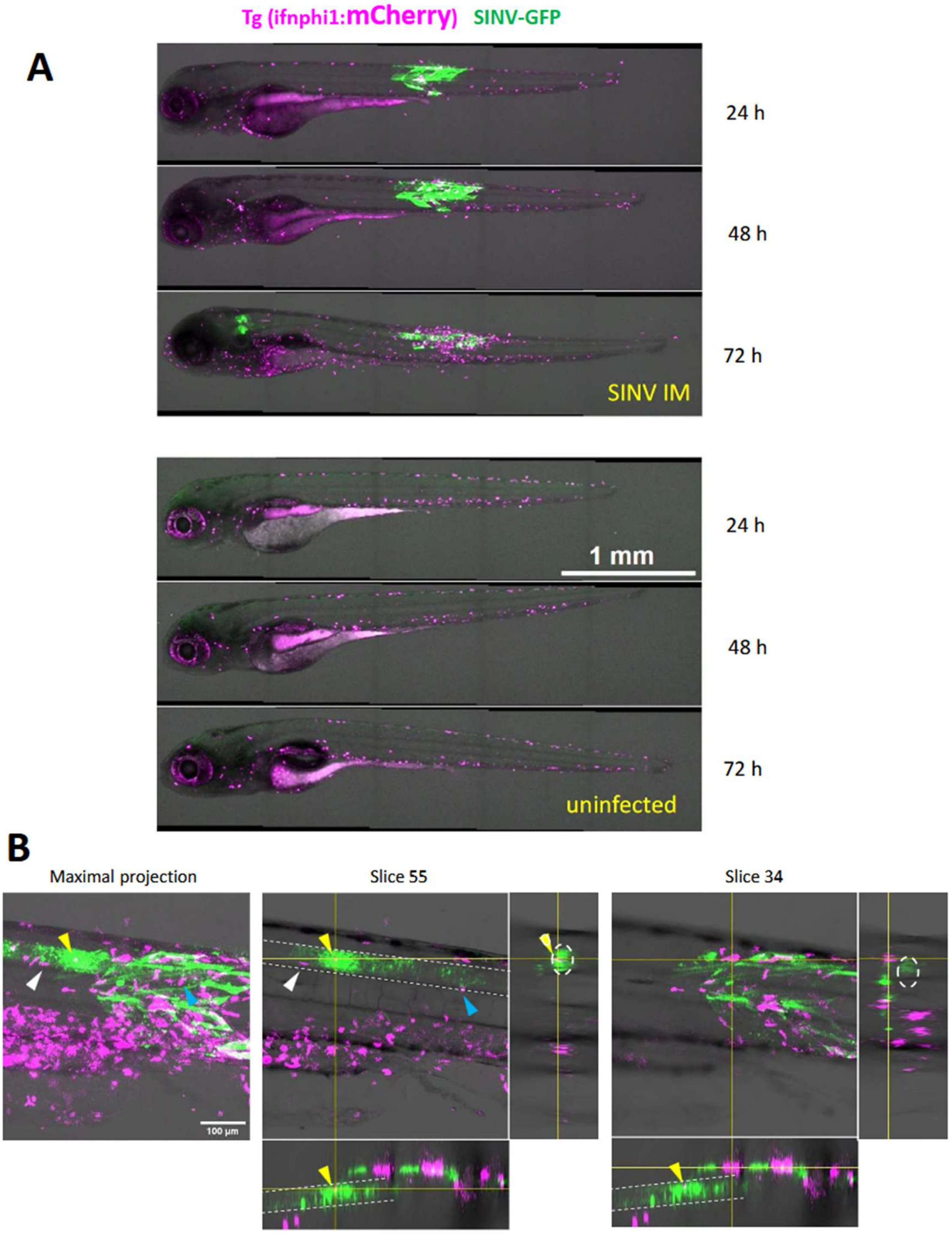

**Figure S4.** IFNphi1 reporter cell imaging during SINV infection. Live confocal imaging, merge of transmitted light and red and green fluorescence channels. A. Whole-body images of one infected and one uninfected representative larva over time, maximal projections. B. Detail of a SINV-infected larvae imaged at 72h post-infection, with maximal projection on the left and two confocal sections with orthogonal projections, one through the spinal cord (slice 55, middle) and one through the overlaying muscle layer (slice 34, right). Yellow arrowhead points to a mCherry positive cells inside the spinal cord, surrounded by infected neurons; white arrowhead points to another mCherry positive cell located just under the envelope the spinal cord, and blue arrowhead to a cell that appears to be entering the spinal cord. White dotted lines indicate the boundary of the spinal cord.

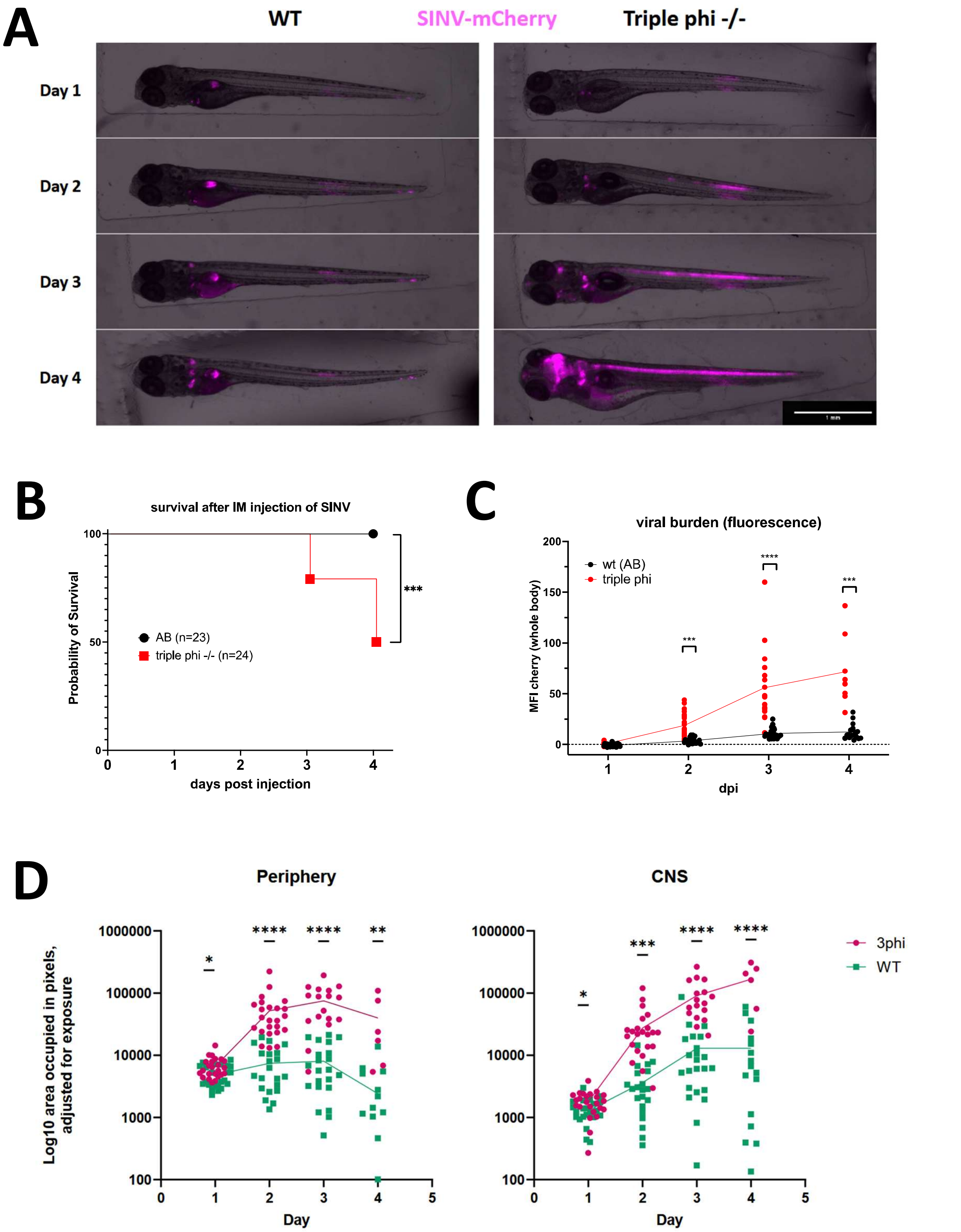

**Figure S5.** IFN-deficient larvae are highly susceptible to SINV infection. Wild-type and homozygous triple phi mutants were injected IM with SINV-mCherry and followed until 4dpi. A. representative outcome of the infection in individual larvae, merge of transmitted and fluorescence images, adjusted for comparable exposures. B. survival of larvae in that experiment. C. Global assessment of viral burden by measurement of mCherry fluorescence on the whole larva. D. decomposition of mCherry fluorescence in the peripheral tissues and CNS areas using the cellprofiler pipeline.

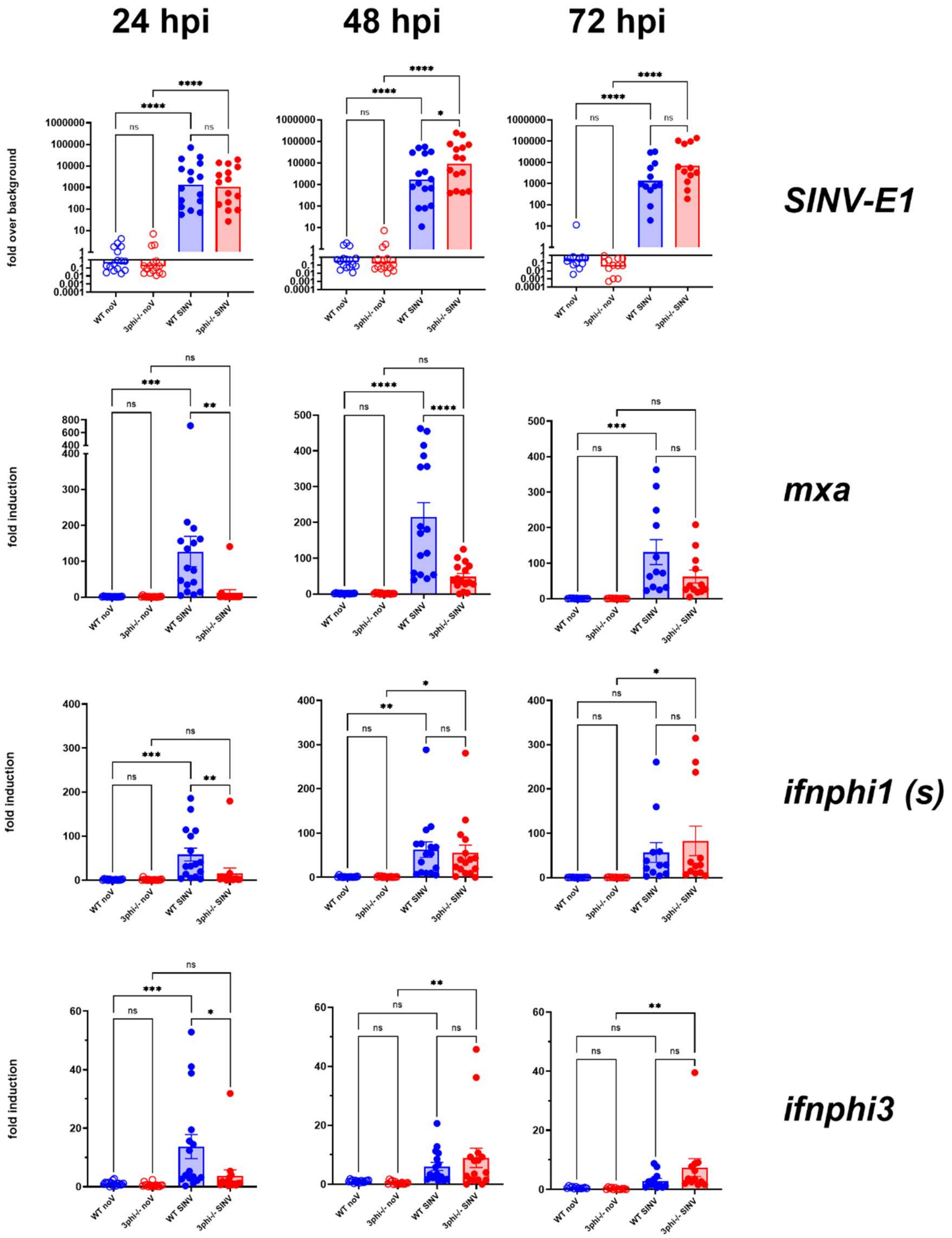

**Figure S6.** qRT-PCR measurement of infection and host response in wild-type and triple-IFN-deficient mutants. Pool of three independent experiments. Values are expressed relative to the mean of WT control samples. Statistical analysis conducted by lognormal ordinary one-way ANOVA.

### Supplementary text 1: Estimation of occurrences of successful CNS invasion events

The number of separate events that resulted in successful invasion of the CNS by SINV was estimated under the assumption that these events were random and independent of each other. According to Poisson's distribution, if a random event happens in average $\lambda$ times per trial over a large number of trials, then the probability of that event not happening at all for any single trial is $e^{-\lambda}$. Considering that the GFP and mCherry viruses are equally invasive, if CNS invasion occurs in average $\lambda$ times per larva, then it will occur $\lambda/2$ times for one given color, and absence of invasion by that color has a probability of $e^{-\lambda/2}$ per larva. Thus, the probability of no invasion at all is $e^{-\lambda}$, that of invasion with any single color would be $2*(e^{-\lambda} - e^{-\lambda/2})$, and that of two-color invasion is $1 - e^{-\lambda} - 2(e^{-\lambda} - e^{-\lambda/2})$.

We deduced from these probabilities the expected occurrences of two-color invasion, single-color invasion, and lack of invasion for a large number of larvae (n=1000) with a range of values of $\lambda$ from 0.1 to 5 (with 0.1 increments) (Annex 1 figure, left panel).

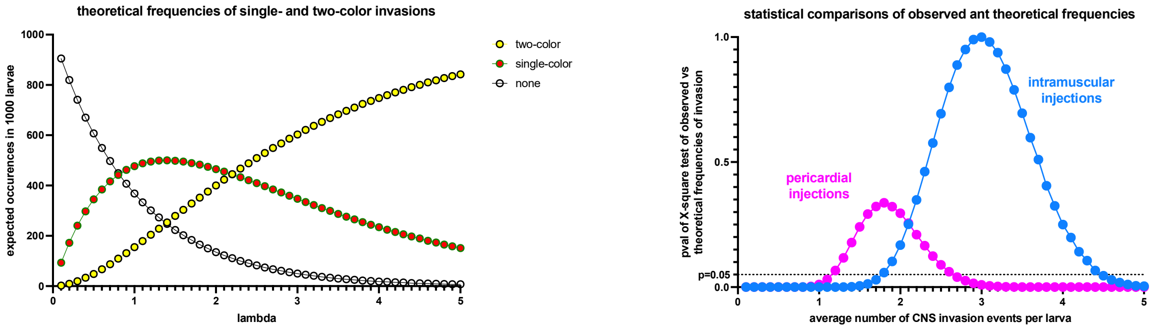

We then compared these frequencies with our observational data using $\chi$-square tests. The range of likely values of $\lambda$ was that for which the observational and theoretical values were not statistically different ($p>0.05$). We thus plotted the pValue of a comparison of the theoretical and observed values of $\lambda$ from 0 to 5. (Annex 1 figure, right panel). Thus, for pericardial injection, the most likely value is 1.8 (range 1.2-2.6) and for intramuscular injections the most likely value is 3.0 (range 1.7-4.4).

**Supplementary text2: Construction of a mathematical model of SINV infection**

We first built a single-compartment model of the infection for the periphery, with the help of the Berkeley Madonna software (Annex Figure 1). The model, essentially based on (Best et al., 2017), allows for a variety of actions of the interferon response, so they can be tested in turn.

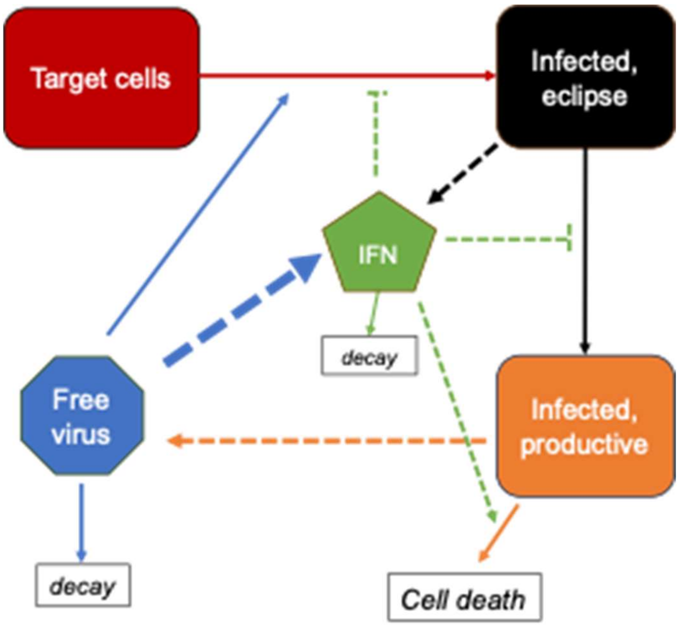

**Annex Figure 1.** Flowchart of the single-compartment model.

The model starts with a population of $10^5$ target cells (T). In presence of free virus, some of these cells get infected, entering first an eclipse phase (E) during which they do not produce virus, before transiting to the productive phase of infection (P), where they start to release new infective virions at a fixed rate. We neglect death or proliferation of target or eclipse cells, but the cytopathic effect (CPE) causes death of productively infected cells at a given rate. Virions decay at a fixed rate and are also lost when infecting a target cell. In absence of a host response, this model is governed by the following equations:

| | |
|---|---|
| $dT/dt = -\beta V(T/T_0)$ | where $\beta$ is the rate of infection ($T_0$ is the initial population) |
| $dE/dt = \beta V(T/T_0) - kE$ | where k is the rate of transition from eclipse to productive |
| $dP/dt = kE - \delta P$ | where $\delta$ is the rate of CPE-induced death |
| $dV/dt = \gamma P - cV - \beta V(T/T_0)$ | where $\gamma$ is the rate of virions production per productively infected cells; and c is the rate of virus decay |

By fluorescence microscopy, we detect the first wave of infected cells in periphery around 7 hours post inoculation; as the fluorescent reporter is co-expressed with structural genes, this corresponds to entry into productive state. Protocols used to titrate SINV typically start with a 1- or 1.5-hour adsorption phase.  Therefore, we set the infection rate of target cells by free virus (β) at 0.5 per hour, implying a median time of 1 hour for a virus to cause a cell to enter eclipse phase, and the transit rate from eclipse to productive state (k) at 0.1 per hour, corresponding a median duration of the eclipse phase of ~6 hours.

The death rate of productively infected cells (δ**)** was set at 0.03 per hour, corresponding to a median survival time of ~24 hours of productively infected cells, a value consistent with our imaging data. Half-life of SINV in culture medium is 4 hours at 37°C (Purifoy et al., 1968); it should be a bit longer at 28°C, so we set virus decay rate (**c**) at 0.1 per hour (i.e. half-life of 6 hours).

After setting these parameters at plausible values, we were left with the more difficult task of estimating the rate of virus production per productively infected cell (γ). However, we know that larvae lacking IFN response die from overwhelming infection between 48 and 72 hours after inoculation of ~30PFU of SINV (Figure 7 of main paper). We thus ran our model with an initial value of 30 for **V** and various values of γ (Annex Figure 2). Except for extremely low values of γ (<0.036) where infected cells may die before producing new virus, this differential equation system ultimately leads to complete infection. We considered that larva death would occur when half the cells would have become productively infected or dead (e.g, $T+E<T_0/2$) (because flow cytometry analysis of the most heavily infected larvae resulted in at most 40% virus-fluorescent cells; see Figure S3 in the main paper). To our surprise, this threshold was reached between 48 to 72 hours for values of γ ranging from 0.6 and 1.5, much lower than we would have anticipated. Thus, according to our model and assumptions, productively infected cells release only one new infective viral particle every hour.

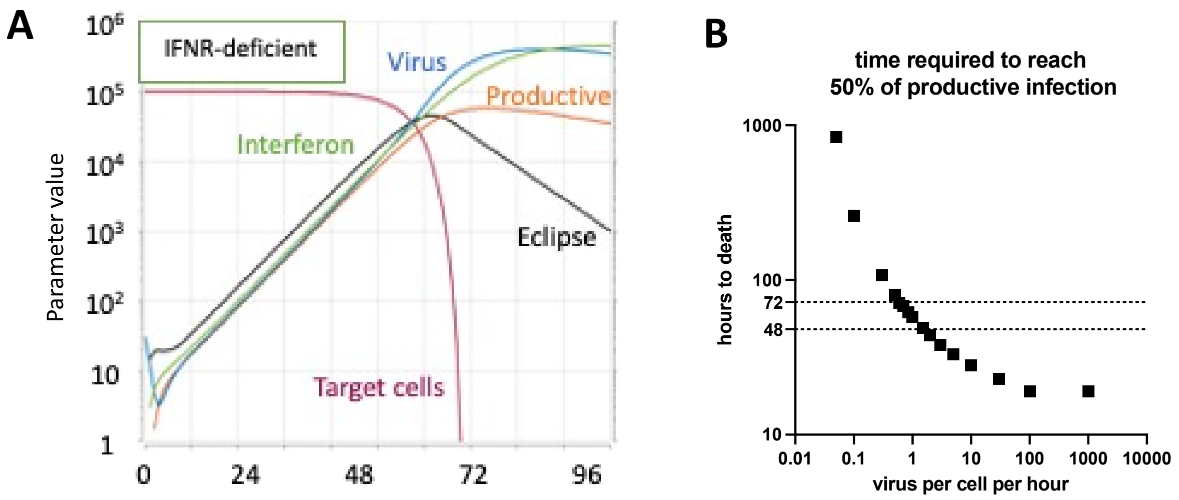

**Annex Figure 2**. Results of running the model without an interferon response. A, simulation of model evolution with a virus production rate (γ) set a 1 per infected cell per hour. Other parameters: β=0.5, δ=0.03, k=0.1, c=0.1. B. Effect of varying the value of γ on the time required for half of the initial cell population to have died or entered productive phase.

We tested if this result was highly sensitive to the previous choice of parameters, by doubling and halving them one by one while keeping $\gamma=1$. The highest sensitivity was observed for the rate of transit from eclipse to productive state **k** (Annex Figure 3A). If this rate was halved (correspond to a mean transition time of 14 hours), the threshold of 50% infected cells would be reached for values of γ ranging from 1.3 to 4 (Annex Figure 3B). Thus, even for this low value of transit mean time, the rate of infectious virus production would be no more than a few virions per hour for each productively infected cells.

For the rest of the modelling, we kept k=0.1 and γ=1 (resulting in the death of IFNR-deficient fish in 58 hours).

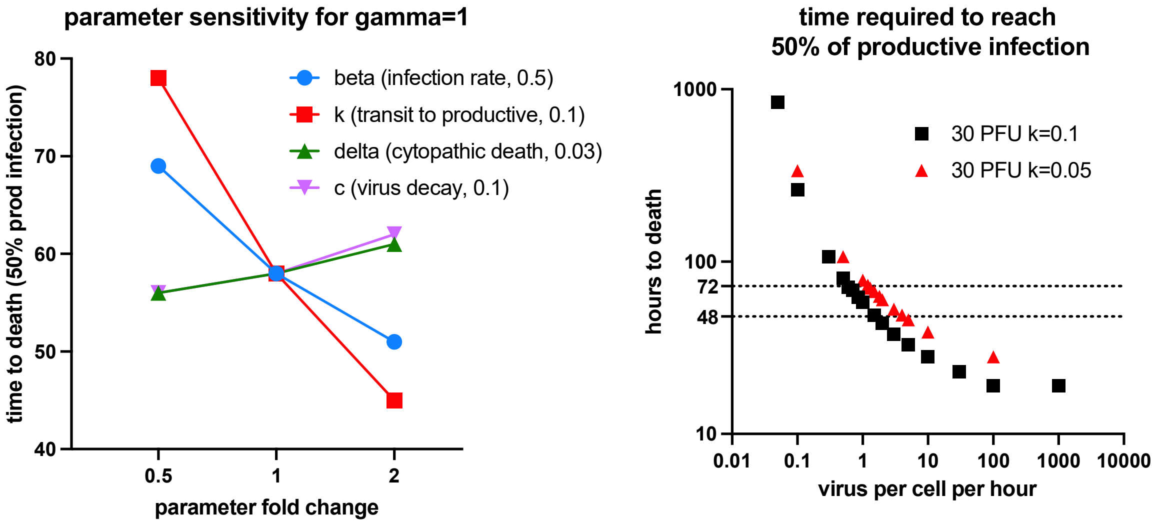

**Annex Figure 3**. Parameter sensitivity analysis A. sensitivity of the model to initial parameter choice. B. effect of variation of the virus production rate for two different values of transit rate.

We then incorporated the role of the interferon response, adding as a last variable the amount of free interferon (**F**), which starts at zero. We considered two detections pathways for the virus: either cytosolic viral RNA (RLR pathway) or extracellular/endosomal virions or fragments thereof (TLR pathway). RLR pathway would be activated in infected cells, but since SINV, like many viruses, shuts off host cell translation, we consider that no IFN would be released by productively infected cells; only cells in eclipse phase would release IFN via this pathway. By contrast, we hypothesize virions would be detected by sentinel cells that do not get infected, as is the case for macrophages (Passoni et al., 2017). Thus, RLR production is proportional to the value of E and TLR production to the value of V. In addition, IFN decays at a constant rate, leading to this equation:

$dF/dT = \rho E + \tau V - \eta F$ $\rho$ is the rate of IFN release by eclipse phase cells

$\tau$ is the rate of IFN released upon detection of virions

$\eta$ is the rate of IFN decay

In humans, half-life of recombinant IFN$\alpha$ has been measured around 6 hours (Gutterman et al., 1982). We thus set $\eta$ at 0.08 for a half-life of zebrafish IFN around 8 hours at 28°C.

We could not guesstimate values for $\rho$ and $\tau$, but we expect that both pathways contribute to the IFN pool based on our findings where knock-down of the key RLR pathway adaptor MAVS reduced but did not abolish the IFN response of zebrafish larvae infected with the closely related chikungunya virus (Palha et al., 2013). Thus we started by setting them at an equal value of 0.1 each (note that the value of the IFN unit is arbitrary – this choice will result in values of F of similar magnitude as V, which is convenient for the graphical representation). If IFN is inactive, this results in an exponential increase of E, P, V and F during two days, until almost all target cells have been infected (Annex Figure 2A).

We then considered that IFN may counteract the infection via three mechanisms: preventing infection of target cells (if a virus attempts to infect a target cell but fails due to IFN response, it is still lost from the free virion pool), preventing transit of eclipse cells into the productive state, and accelerating the death of productively infecting cells (by any mechanism, such as direct effect of IFN on these cells or by activating killer immune cells). Following (Best et al., 2017), we assumed that IFN inhibited these events in a near-linear fashion, thus modifying our initial equations by adding the terms in bold below:

$dT/dt = (-\beta V(T/T_0))\mathbf{/(1+\varepsilon F)}$ $\varepsilon$ is the inhibition of infection by IFN

$dE/dt = \beta V(T/T_0)\mathbf{/(1+\varepsilon F)} - kE\mathbf{/(1+\phi F)}$ $\phi$ is the inhibition of transit by IFN

$dP/dt = kE\mathbf{/(1+\phi F)} - \delta P\mathbf{*(1+\theta F)}$ $\theta$ f is the induction of death by IFN

$dV/dt = \gamma P - cV - \beta V(T/T_0)$

Our experimental results indicate that in IFN-competent larvae, the amount of productively infected cells in periphery should peak at approximatively 36 hours, then decline. We thus assessed time and height of the peak values of P to fit the three unknown IFN-based inhibition parameters.

We first tested equal values for ε, ϕ and θ on the model (Annex Figure 4). Except at very low values ($10^{-5}$ or less) for which all target cells got infected, adding this IFN response resulted in a stabilization (but not disappearance) of the infection after a peak, which was reached earlier if the IFN response was stronger. The height of this peak was inversely proportional to the strength of the IFN response. The peak was reached at ~36 hours for $\varepsilon = \phi = \theta = 0.003$, with 230 infected cells, a value that appears consistent with our microscopic observations. About $10^3$ cells enter eclipse phase, and the decline in target cell numbers is imperceptible.

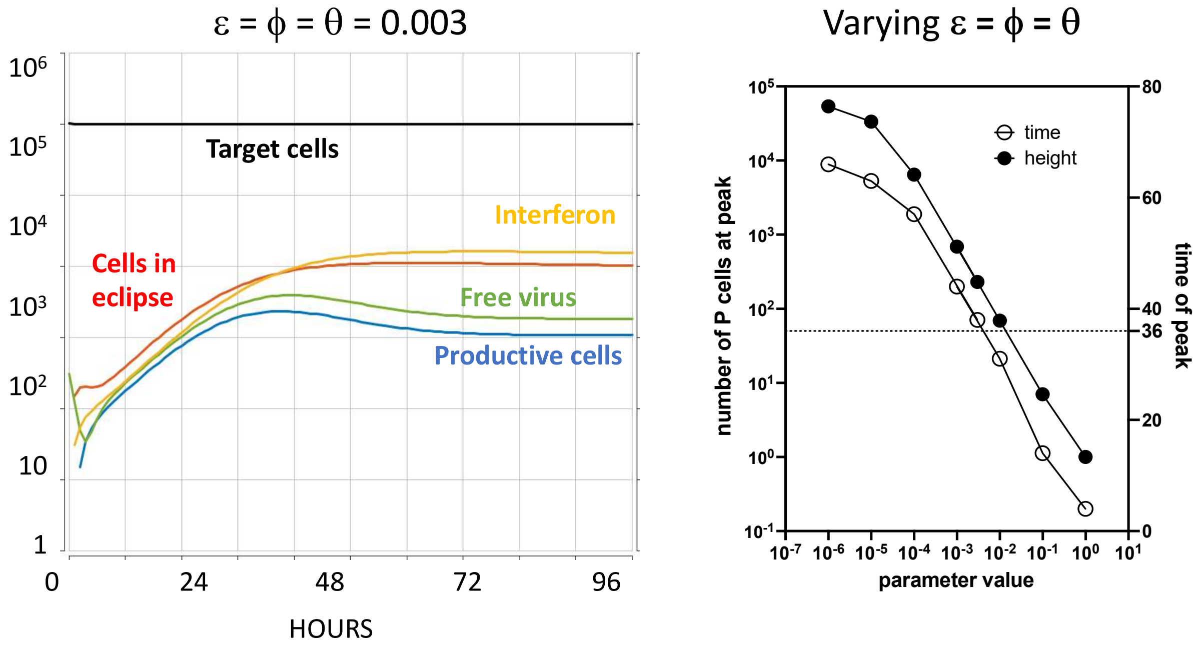

**Annex Figure 4.** initial manual fitting of the IFN inhibition values. Parameter values for the simulation on the left: β=0.5, δ=0.03, k=0.1, c=0.1, γ=1, ρ=0.1, τ=0.1, η=0.08, $\varepsilon = \phi = \theta = 0.03$.

To compare the relative importance of these three modes of inhibition, we first canceled each of them in turn, keeping the other two at 0.03 (Annex Figure 5A). In simulations where death of productive cells was not accelerated by IFN, no peak of infection is observed, only a plateau, this this mode seems critical to reflect our experimental data. The importance of the other two modes is less clear. If transit is not blocked, the peak is attenuated; if infection is not blocked, a large population of eclipse cells is retained. In in our model, not only was accelerated death necessary, it was also sufficient; with a value of θ set at 0.01, our simulation reached a peak at 36 hours with dynamics globally similar to that shown of fig 5A.

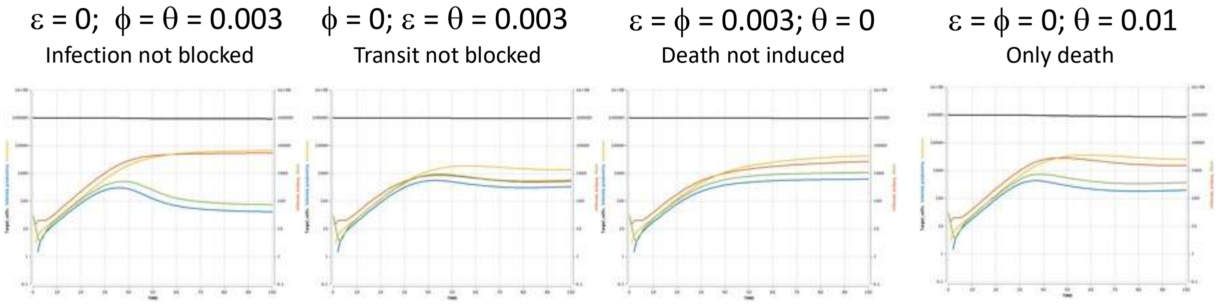

Annex Figure 5. testing the different pathways by which IFN may counteract the infection. Other parameter values: $\beta$=0.5, $\delta$=0.03, k=0.1, c=0.1, $\gamma$=1, $\rho$=0.1, $\tau$=0.1, $\eta$=0.08.

Finally, we tested the exclusion the RLR or the TLR pathways for IFN induction. Keeping $\varepsilon = \phi = \theta = 0.003$, we set either r or t to 0 (Annex Figure 6). Interesting, RLR detection alone resulted in a clear peak, while TLR detection only resulted in a plateau, indicating that RLR detection was critical while the TLR pathway may not play a major role.

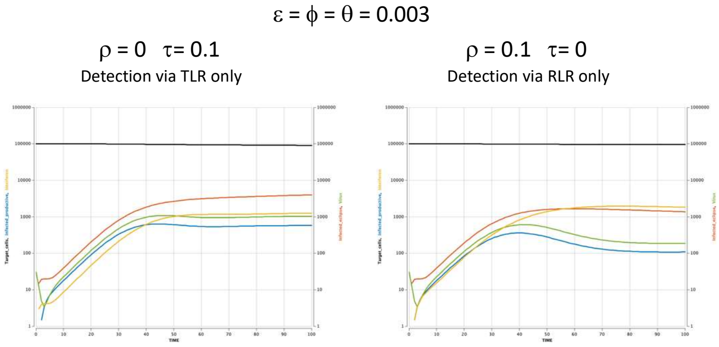

Annex Figure 6. testing the impact of the RLR and TLR pathways. Other parameter values: $\beta$=0.5, $\delta$=0.03, k=0.1, c=0.1, $\gamma$=1, $\varepsilon = \phi = \theta = 0.03$.

Finally, we used Berkeley Madonna's parameter fitting function to find parameters that would best match our actual data. We used our imaging-based quantifications of infection over time in 48 larvae (Figure 5G of main article); the fitted variable was the number of productively infected cells. We ran multi-parameter fitting of 7 parameters: the 5 IFN response-related parameters ($\rho$, $\tau$, $\eta$, $\varepsilon$, $\phi$, $\theta$) but also the CPE-induced cell death rate ($\delta$) and the virus production rate ($\gamma$). Results are displayed on Annex Figure 7. These results suggest that in periphery, IFN is mainly produced by eclipse phase cells ($\rho$>>$\tau$) and IFN has multiple effects, but transit block is dominant ($\phi > \theta > \varepsilon$).

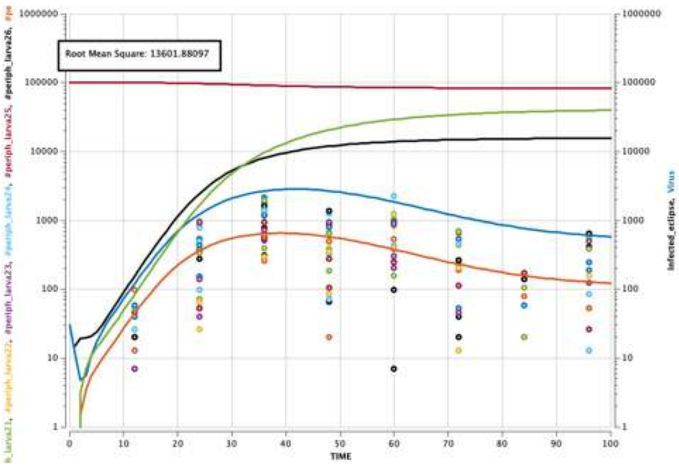

| Parameter | symbol | Before fitting | After multi fitting |
|---|---|---|---|
| infection rate | β | 0.5 | 0.5 |
| transition rate | k | 0.1 | 0.1 |
| CPE death rate | δ | 0.03 | 0.00318 |
| virus production | γ | 1 | 2.39124 |
| virus decay | c | 0.1 | 0.1 |
| IFN release by eclipse cells | ρ | 0.1 | 0.20849 |
| IFN release by sentinel cells | τ | 0.1 | 8.4e-16 |
| IFN decay | η | 0.08 | 0.08 |
| infection block by IFN | ε | 0.003 | 1.8e-4 |
| transit block by IFN | φ | 0.003 | 0.00329 |
| death induction by IFN | θ | 0.003 | 8.2e-4 |
| RMSq | | 16763 | 13601 |

**Annex Figure 7.** Computer-aided fitting of multiple parameters onto measured infection measured in periphery.

Having established parameter values for which our model reflected relatively well SINV infection in the periphery, we turned to the CNS compartment.

First, we assume that the BBB prevents virion and IFN exchange between periphery and CNS so that simulations are run independently in the two compartments. A few neurons initially get infected by axonal transport, and our experimental data indicate, on average, three independent events, with the first neurons expressing virus-encoded fluorescence around 24 hpi. Thus, the simulation starts with no free virus, but 3 cells entering eclipse phase at 12 hpi (corresponding to t=0 here).

First, we manually adjusted the periphery-related parameters guided by literature and experimental data.  Obviously the model parameters for CNS could not be identical to those we determined in periphery, as this would result in rapidly controlled infection.

We considered that the following parameters should remain unchanged between periphery and CNS: rate of infection (β=0.5), rate of transit from eclipse to productive (k=0.1), rate of new virion production (γ=1), rate of virus decay (c=0.1), rate of IFN decay (η=0.08).

Death of infected neurons was a rare event in our live imaging data (except for the very small population of DRG neurons). Accordingly, we reduced the cytopathic effect-induced death rate (δ=0.01 instead of 0.03). This alone had little effect on the simulation.

Neurons are known to be poor IFN producers (Viengkhou and Hofer, 2023). For this reason we strongly reduced the rate of IFN produced by eclipse cells via the RLR pathway (ρ=0.01 instead of 0.1). By contrast, since we observed ifnphi1-expressing leukocytes inside the infected CNS, the sentinel cell TLR pathway IFN source was unchanged (τ=0.1). This still resulted in a clearly controlled infection, but with a 3-fold increase in the maximal number of productively infected cells.

Finally, neurons are also known to have a narrow ISG response to IFNs (Viengkhou and Hofer, 2023). This is consistent with the lack of expression of the MXA:mCherry transgene that we observe in neurons. At this stage we do not know what pathways are affected at this stage, we first tested lowering equally all IFN impact modelled here: blockade of infection, blockade of transition to productive phase, acceleration of death of infected cells. Reducing these effects resulted in a proportional increase of the peak of number of productively infected cells. We observe a strong variability in the CNS of our experimental fish: some appear to control the infection, while others do not. This variance cannot be replicated in our mathematical model which is deterministic; however, parameters yielding an infection that is controlled only very close to the upper limit would reflect this, as small changes would result in this life-or-death outcome. This is attained when the three parameters that reflect IFN control ($\varepsilon$, $\phi$, $\theta$) are reduced 10-fold (Annex Figure 8A). We then tested the relative importance of these three parameters by setting each to zero in turn. The impact on the total numbers of productively infected cells (our observable value) was relatively modest, but dramatic differences were observed in the number of uninfected target cells and eclipse phase cells (Annex Figure 8B).

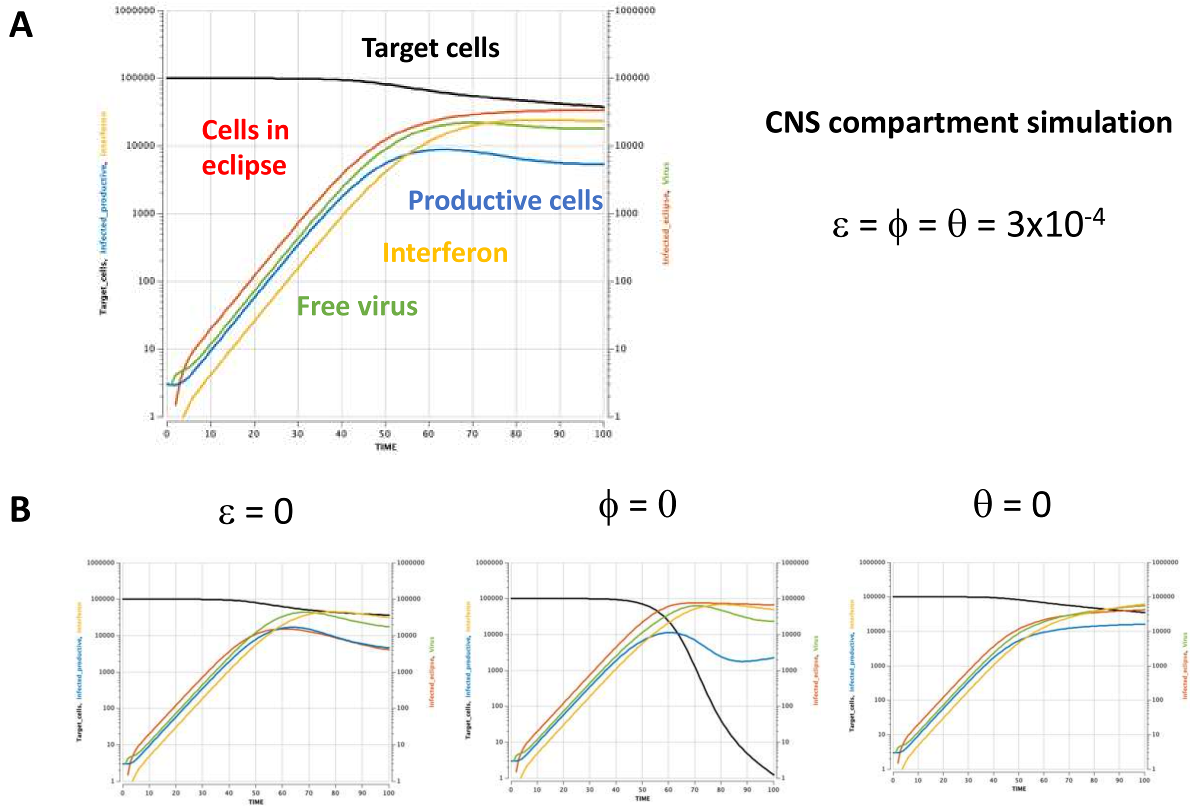

**Annex Figure 8.** Manual daptation of the model to the CNS compartment. A. simulation with all three IFN-impact parameters. Other parameter values: $\beta$=0.5, $\delta$=0.01, k=0.1, c=0.1, $\gamma$=1, $\rho$=0.01, $\tau$=0.1, $\eta$=0.08. B. Model ran with on of the three IFN-impact parameter set to zero, while the other two are set at 3.10e-4. Other parameters as in A.

Finally, we conducted computer-aided multi-parameter fitting based on the actual image-measured values (Figure 5FG of main paper). Interestingly, these simulations, as show on Annex Figure 9, confirm our intuition that sentinel cells produce more IFN than eclipse cells

($\tau >> \rho$). Block transit from eclipse to productive is the only significant action of IFN ($\phi >> \theta > \varepsilon$). The simulation also shows that, unlike in periphery, all neurons ultimately get infected, although most of them stay in eclipse phase.

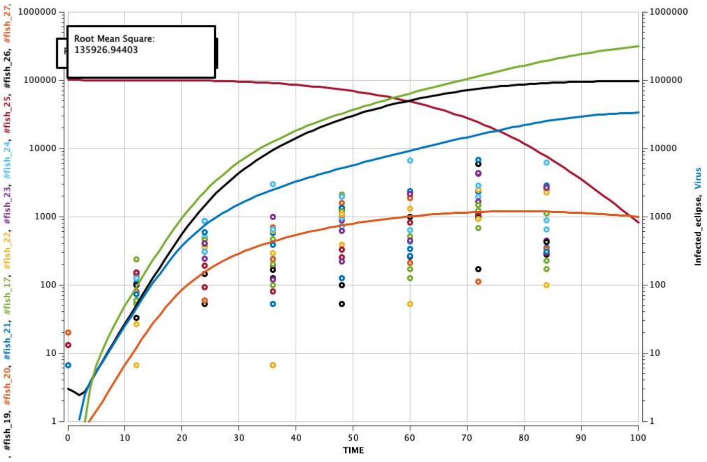

| Parameter | symbol | Initial value | After multi fit |
|---|---|---|---|
| infection rate | $\beta$ | 0.5 | 0.5 |
| transition rate | k | 0.1 | 0.1 |
| CPE death rate | $\delta$ | 0.03 | 0.03 |
| virus production | $\gamma$ | 1 | 3.6 |
| virus decay | c | 0.1 | 0.1 |
| IFN release by eclipse cells | $\rho$ | 0.1 | 1 e-14 |
| IFN release by sentinel cells | $\tau$ | 0.1 | 0.92 |
| IFN decay | $\eta$ | 0.08 | 0.08 |
| infection block by IFN | $\varepsilon$ | 0.003 | 6 e-7 |
| transit block by IFN | $\phi$ | 0.003 | 0.0024 |
| death induction by IFN | $\theta$ | 0.003 | 6 e-6 |
| RMSq | | 135926 | 135926 |

Annex Figure 9. Computer-aided fitting of multiple parameters onto measured infection measured in CNS.

**Supplementary movies**

All movies except S11 & 12 have been obtained by live imaging

**Movie S1 – progression of SINV infection during the first day, whole-body view**
Relative to Figure 2A (top panels)
Tg(elavl3:GFP) zebrafish larva inoculated IM with SINV-mCherry
Confocal timelapse (10x objective), maximal projection
Merge of transmitted light (gray) and fluorescence channels (GFP in green, mCherry in magenta)
Imaged from 6.6 to 24 hpi every 76 minutes

**Movie S2 – progression of SINV infection during the second day, tail view**
Relative to Figure 2B
Tg(vsx2:GFP) zebrafish larva inoculated IM with SINV-mCherry
Confocal timelapse (10x objective),, maximal projection
Merge of transmitted light (gray) and fluorescence channels (GFP in green, mCherry in magenta)
Imaged from 24 to 40 hpi every 95 minutes

**Movie S3 – progression of SINV infection during the second day, whole-body view**
Relative to Figure 2A (bottom panels)
Tg(elavl3:GFP) zebrafish larva inoculated IM with SINV-mCherry
Confocal timelapse (10x objective), maximal projection
Merge of transmitted light (gray) and fluorescence channels (GFP in green, mCherry in magenta)
Imaged from 32hpi to 40 hpi every 76 minutes

**Movie S4 – emergence of the first infected neuron, tail view**
Relative to Figure 2C
Tg(mnx1:GFP) zebrafish larva inoculated IM with SINV-mCherry
Confocal timelapse (10x objective), maximal projection
Merge of transmitted light (gray) and fluorescence channels (GFP in green, mCherry in magenta)
Imaged from 24hpi to 35nhpi every 72 minutes

**Movie S5 – 3D rendering of the area around the first infected neuron of movie S4**
Tg(mnx1:GFP) zebrafish larva inoculated IM with SINV-mCherry at 35 hpi
Confocal imaging (10x objective), 3D projection around the Y-axis
Merge of fluorescence channels (GFP in green, mCherry in magenta)

**Movie S6 – infection of a DRG sensory neuron**
Relative to Figure 2D
Tg(ngn1:GFP) zebrafish larva inoculated IM with SINV-mCherry
Confocal timelapse (40x objective), deconvolution and maximal projection
Merge of transmitted light (gray) and fluorescence channels (GFP in green, mCherry in magenta)
Imaged from 28 to 43 hpi every 34 minutes

**Movie S7 – Early infection of a motoneuron, followed by DRG neurons**
Relative to Figure 2E
Tg(ngn1:GFP) zebrafish larva inoculated IM with SINV-mCherry
Confocal timelapse (40x objective), deconvolution and maximal projection
Merge of fluorescence channels (GFP in green, mCherry in magenta)
Imaged from 28 to 43 hpi every 34 minutes

**Movie S8 – advancing infection in the spinal cord**
Relative to Figure 3A, and to movie S7
Tg(ngn1:GFP) zebrafish larva inoculated IM with SINV-mCherry
Confocal timelapse (40x objective), deconvolution and maximal projection
Cropped fluorescence channel for mCherry (in grays)
Imaged from 28 to 43 hpi every 34 minutes

**Movie S9 – death of infected DRG neuron**
Tg(ngn1:GFP) zebrafish larva inoculated IM with SINV-mCherry
Confocal timelapse (40x objective), deconvolution and maximal projection
Merge of fluorescence channels on cropped area (GFP in green, mCherry in magenta)
Imaged from 28 to 43 hpi every 34 minutes

**Movie S10 – response of macrophages to SINV infection**
Relative to Figure 6D
Tg(mfap4:mCherryF) zebrafish larva inoculated IM with SINV-GFP and labelled with NucRed Live
Confocal timelapse (10x objective), deconvolution and maximal projection
Merge of fluorescence channels (GFP in green, mCherry in magenta, NucRed in blue)
Imaged from 50 to 69 hpi every 21.5 min

**Movie S11 – SINV infection of an IFNR-knockdown endothelial reporter larvae**
IFNR-Morpholino (MO)-treated Tg(fli1:GFP) zebrafish larvae IM with SINV-mCherry
Confocal imaging after fixation at 48 hpi, clarification, immunostaining for GFP (green) and mCherry (magenta). Crop of the hindbrain region; the first frame corresponds to the maximal projection of the Z-stack.
